# Boosting performance of generative diffusion model for molecular docking by training on artificial binding pockets

**DOI:** 10.1101/2023.11.22.568238

**Authors:** Taras Voitsitskyi, Volodymyr Bdzhola, Roman Stratiichuk, Ihor Koleiev, Zakhar Ostrovsky, Volodymyr Vozniak, Ivan Khropachov, Pavlo Henitsoi, Leonid Popryho, Roman Zhytar, Semen O Yesylevskyy, Alan Nafiiev, Serhii Starosyla

## Abstract

This study introduces the PocketCFDM generative diffusion model, aimed at improving the prediction of small molecule poses in the protein binding pockets. The model utilizes a novel data augmentation technique, involving the creation of numerous artificial binding pockets that mimic the statistical patterns of non-bond interactions found in actual protein-ligand complexes. An algorithmic method was developed to assess and replicate these interaction patterns in the artificial binding pockets built around small molecule conformers. It is shown that the integration of artificial binding pockets into the training process significantly enhanced the model’s performance. Notably, PocketCFDM surpassed DiffDock in terms of non-bond interaction quality, number of steric clashes, and inference speed. Future developments and optimizations of the model are discussed.

**Availability:** The inference code and final model weights of PocketCFDM are accessible publicly via the GitHub repository: https://github.com/vtarasv/pocket-cfdm.git.

## Introduction

Molecular docking plays a central role in modern computational drug discovery. Until recently docking was the only available method of predicting the poses of small molecules in the binding pockets of target proteins fast enough to be useful in large-throughput virtual screening projects. Despite its ultimate importance, there is a notable stagnation in improving the docking versatility, accuracy, computing cost, and predictive power ^1,2^. Being based on inevitably simplified empirical potentials of intermolecular interactions and lacking explicit solvent the docking scoring functions have arguably reached the plateau of practical accuracy. Despite a number of recent developments in the field, all of them are incremental improvements and domain-specific tuning rather than technological breakthroughs.

However, recent advancements in deep learning methodologies for predicting ligand poses in the protein binding pockets provide a promising alternative to docking algorithms. These techniques can be categorized into two primary groups.

The models from the first group utilize a one-shot inference regression-based approach ^3,4^. They are developed with the aim of very fast inference, which is superior to traditional docking simulations. The Geometric Vector Perceptrons (GVP) and Equivariant Graph Neural Networks (EGNN) are the most popular architectures for those types of models ^5–7^. Such techniques as EquiBind ^4^ and TANKBind ^3^ demonstrated efficacy in predicting protein-ligand binding structure without prior knowledge about the binding pocket (also known as blind docking) while being faster than traditional docking techniques by several orders of magnitude. However, they still suffer from unrealistic ligand conformations and numerous sterical clashes in predicted complexes, which puts them behind the docking approaches in terms of structure quality and reliability. A possible reason for this is a mismatch between the objectives of molecular docking and the regression paradigm. Particularly, the accuracy metrics in molecular docking are based on structural similarity, rather than a regression loss.

The second and currently state-of-the-art approach uses generative AI models, which aim to be on par or better than classic docking techniques in terms of accuracy and structure quality. The DiffDock ^8^, a current leader in the field, demonstrates superior performance in comparison to some conventional docking techniques and the previous ML models in the blind docking scenarios. It produces much less steric clashes than its rivals and generates realistic ligand conformations. However, the enhanced precision of DiffDock comes at the cost of a substantial computational burden, which is on par with that of traditional docking methods. Since there is a significant potential for improvement in terms of inference speed, it makes generative models the most promising in the field at the moment of writing.

The major bottleneck, which hampers further improvement of the generative ligand pose prediction models, is the inherently limited amount of the training data. The overall number of experimentally determined protein-ligand complexes resolved by X-ray, Cryo-EM, or NMR techniques is now below 20,000. The PDBbind database ^9,10^, which is a primary dataset for machine learning in protein binding site prediction ^11,12^ and ligand pose prediction ^3,4,8^, contains 19,443 distinct protein-ligand complexes with the binding activity annotations (version 2020). Out of this, 2,709 entries involve peptide ligands or multiple molecules in a single binding pocket; 3,827 have an insufficient resolution (2.5 angstroms and more); 3,523 contain weak binders (Kd/Ki/IC_50_ > 10 μM) and 216 lack confident activity measurements. Thus there are only 10.270 high-quality complexes, which could be used for model training.

Another issue of the experimental complexes is ligand data sparsity. There are 12,815 distinct small molecule ligands in PDBbind, of which only 1,655 appear in two or more entries (Figure 1). It indicates that ML-based approaches are mostly presented with a particular ligand bound to a single protein without any information about the possible variability of its binding modes. This might lead to overfitting to a single ligand pose, which wouldn’t be the case when working with more dense data as was demonstrated for the affinity prediction models ^13^.

**Figure 1.**
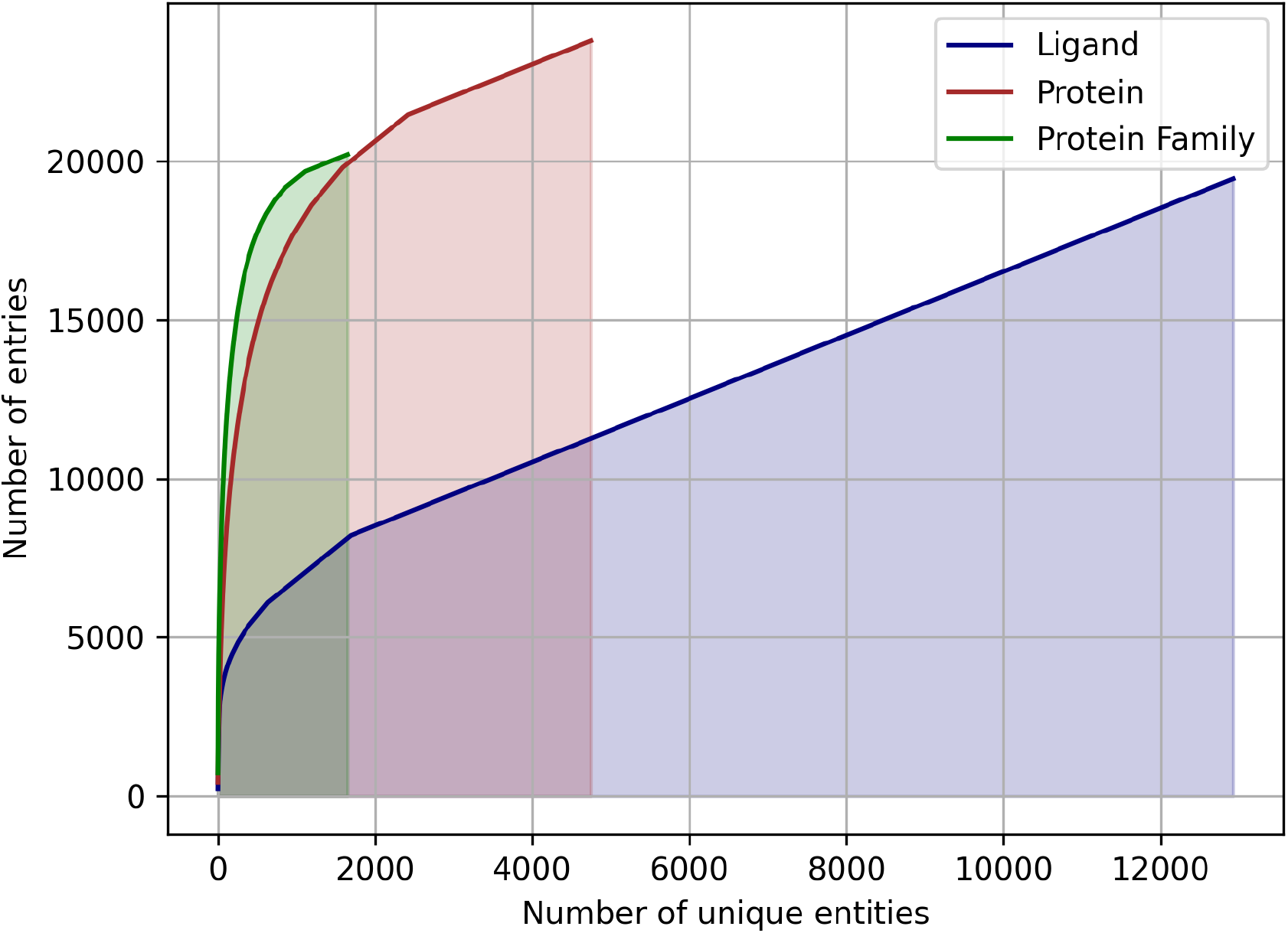
The cumulative sums of entries in the PDBbind database assigned to a unique ligand, unique protein, or unique protein family. Note that the count of proteins and families may exceed the total number of complexes in PDBbind because a single PDB structure may be assigned to multiple entries.

In addition, the proteins are represented very unevenly in available data. The Protein Data Bank (PDB) identifiers of 19,443 protein-ligand complexes are associated with 4.749 unique UniProt^14^ identifiers, but 1.500 most frequently occurring identifiers accounting for ∼80% of the total number of complexes (Figure 1). The same is true for the protein families: a total of 1,646 distinct protein families and superfamilies are present but the top 100 families account for almost 60% of complexes (Figure 1). Thus, despite a significant overall variety of proteins, there is an obvious overrepresentation of some proteins and protein families, which may lead to model overfitting and imbalance.

There is little doubt that an insufficient overall number of samples, limited ligand diversity, and protein representation imbalance in the available data for model training are impairing the accuracy of ML-based approaches to the ligand pose prediction. These limitations are especially noticeable when comparing the amount and quality of the training data with a requirement for the model to operate on arbitrary protein targets and arbitrary ligands from an immense chemical space of at least ^60^ organic molecules ^15^.

It is clear that the dataset of experimentally resolved structures will not grow fast enough to satisfy the demands of the exploding field of ML ligand-protein binding prediction thus other approaches are needed for overcoming the lack of training data.

In this study, we develop an approach of augmenting the training set of the protein-ligand complexes with artificial data, which mimics real protein binding pockets in a number of structural characteristics. The statistical distributions of artificial pockets’ parameters are fitted to the respective distributions of real protein-ligand structures so that both types of data could be used together seamlessly.

The idea of our approach is based on the assumption that the number of favorable interaction geometries between the protein amino acids and the chemical groups of the ligands is finite and is represented sufficiently well in the available experimental data. What is presumably lacking, is the sampling of all possible combinations of such interactions within the binding pocket. In other words, we assume that available experimental structures provide decent statistics about the preferable chemical identity of interacting atom pairs, distances between the atoms, and orientations of the corresponding chemical groups. However, only a very small fraction of all possible combinations of such pairs is observed in real proteins.

If one generates a large number of “artificial binding pockets”, which follow the same statistical distribution of the interacting atom pairs as the real ones, but sample a much larger variety of their combinations, it might be possible to overcome the undersampling and to train the model on a more complete set of data.

Although we do not have a strong independent proof of our hypothesis, we decided to validate it experimentally by developing an algorithm for generating artificial pockets, compiling a dataset consisting of artificial and real pockets, and measuring the performance of the diffusion generative model, which is inspired by DiffDock, and is trained on such augmented data - PocketCFDM (Pocket Conformation Fitting Diffusion Model). We show that PocketCFDM outperforms DiffDock, which is currently the state of the art in the field, in terms of generated ligand poses correctness (less steric clashes and more favorable non-covalent interactions). We also discuss future prospects of our methodology in terms of improving its predictive power and the speed of inference.

## Methods

### Protein and ligand preprocessing

The Python API of the RDKit v. 2021.03 was utilized for loading, processing, and feature generation of small molecules. A custom protein processing module was developed to extract protein data from the PDB files and to generate the necessary features for model training. This module utilizes the PDB atom names to obtain atom-level graph features, rather than relying on a third-party software to infer them. This approach decreases the exclusion rate for processed proteins due to inevitable inconsistencies in the PDB files.

In order to assess the protein-ligand interactions, we employed the SMILES arbitrary target specification (SMARTS) substructure search to classify the ligand atoms or chemical groups into the following categories: hydrophobic, aromatic, amide, donor, acceptor, cation, anion, or halogen. The protein atom assignment was conducted using a predefined mapping of the standard PDB atom names (Table S1).

Prior to the assessment of non-covalent interactions, both proteins and ligands were protonated using the <u>Reduce</u> software in order to account for hydrogen bonding correctly. Additionally, we considered possible alternative positions of hydrogens, such as within hydroxyl or amine groups. The source code of the preprocessing module is available: https://github.com/vtarasv/rai-chem.git.

### The choice of non-bond interactions

The hydrophobic and electrostatic interactions as well as the hydrogen bonding (favorable non-bond interactions) were assessed between the protein and the ligand. We also accounted for unfavorable interactions, such as donor-donor atom pairs in close proximity, to improve the overall quality of artificial binding pockets by omitting such interactions. The summary of all used non-bond interactions is shown in Table S4. The choice of included interactions is based on a compromise between the multiple approaches in the literature ^16–19^, commonly used cheminformatics software ^20–22^ and widely-used molecular modeling tool <u>BIOVIA Discovery Studio Visualizer</u>.

### Ligand-protein interaction statistics

The protein-ligand complexes with known 3D structures were taken from the PDBbind dataset ^9,10^. Only the entries that satisfy the following criteria were used: the presence of a single small molecule ligand, resolution below 2.5Å, and an activity/affinity less than 10 μM. A total of 10.270 protein-ligand complexes were selected. Among them 1.805 ligand files were found to be unreadable, resulting in a final count of 8.465 complexes that were used in this work.

The following statistical information was extracted:

- The probability of a specific ligand substructure (particular atom type, aromatic ring, or amide group) to participate in a protein-ligand interaction.
- The distribution of the number of the binding pocket substructures that are connected to ligand substructures through a specific interaction type.
- The distribution of amino acids involved in particular interaction types.
- The distributions of distances and angles involved in particular interaction types.

### Artificial pockets generation

We utilized the PeptideBuilder package^23^ to produce a collection of 20 amino acids in the PDB format, as well as 400 dipeptides representing each possible permutation of two amino acids. The amino acids and dipeptides were flanked by GLY residues on both sides and served as the basic building blocks for artificial pockets. The inclusion of flanking GLY residues helps in the generation of the correct peptide conformers, which are restrained by the peptide bonds on each side of the building blocks.

Artificial pocket generation is an iterative process of placing the building blocks around a small molecule in order to form a realistic network of non-covalent interactions. A total of 13 potential interactions, both directed and undirected, were taken into account during the pocket construction (Table 1):

**Table 1.** Non-bonded interactions taken into account during artificial pocket construction.

| # | Pocket feature | Ligand feature | Interaction type |
| --- | --- | --- | --- |
| 1 | Aromatic ring | Aromatic ring | pi stacking |
| 2 | Amide group | Aromatic ring | amide-pi |
| 3 | Aromatic ring | Amide group | amide-pi |
| 4 | Aromatic ring | Cationic atom | cation-pi |
| 5 | Hydrogen bond donor | Hydrogen bond acceptor | hydrogen bond |
| 6 | Hydrogen bond acceptor | Hydrogen bond donor | hydrogen bond |
| 7 | Hydrogen bond acceptor | Halogen atom | halogen bond |
| 8 | Cationic atom | Anionic atom | electrostatic |
| 9 | Anionic atom | Cationic atom | electrostatic |
| 10 | Cationic atom | Aromatic ring | cation-pi |
| 11 | C or S atom | F atom | hydrophobic |
| 12 | C or S atom | Cl, Br or I atom | hydrophobic |
| 13 | C or S atom | C or S atom | hydrophobic |

The pocket construction starts from the particular small molecule conformer (referred as ligand hereafter). For each of the 13 interactions listed in Table 1, the following steps are performed:

1. Find all the features of ligand *L*, which are compatible with the current interaction type *i.* Each such feature is denoted as *F_Li_*.
2. Given experimental probability *P* of finding *F_Li_* among all interactions of type *i* and the random number *p*, determine whether the new interaction should be added if p<P.
3. Select the peptide building block *B* to be placed by taking all building blocks with the features compatible with *i* and randomly selecting one of them according to the experimentally determined probability of the corresponding residue to participate in the interaction *i*.
4. For the chosen building block *B,* generate a random conformer taking into account the peptide bonds to flanking GLY residues. Then delete the flanking GLY residues.
5. Randomly sample the distance *d* between the *F_Li_* and the matching feature of *B* from experimentally determined distributions and place the building block at a determined location.
6. Repeat the following steps until no steric clashes or unfavorable interactions are found between the building block and the ligand and between the building blocks:

a. Randomly rotate and translate the building block *B* preserving the distance *d*.
b. Determine whether the angular criteria of interaction *i* are met, if applicable.
c. If the maximal number of tries (2000 by default) is reached, the building block is skipped.

Using this algorithm 8.465 artificial pockets were generated (one for each ligand from the PDBbind database). The distributions of the non-bond interactions of the generated pockets were computed and compared to the experimental distributions. Due to a good match of the distributions, no further tuning of the algorithm was required.

The examples of randomly selected artificial pockets can be found in the Supporting Information (Figure S1).

### Model training and testing datasets

In order to cover the maximal diversity of the ligands we employed the ZINC20 database of commercially available chemicals widely used for virtual screening ^24,25^. The “In-Stock’’ category of chemicals was chosen, resulting in a collection of 13 million molecules represented by SMILES. The compounds were standardized using the ChEMBL Structure Pipeline ^26^ (https://github.com/chembl/ChEMBL_Structure_Pipeline). Particularly, we eliminated duplicates, compounds with less than seven heavy atoms, and large molecules with a molecular weight exceeding 750 g/mol. This resulted in 9,041,707 compounds. A subset of compounds (size depends on the model training settings) was randomly selected and used for artificial pockets generation. The pocket generation was repeated for each model training epoch.

In addition to the artificial pockets, the real binding pockets from the PDBbind database were added to the training set. These were defined as the residues with at least one heavy atom within a 5Å from any heavy atom of the ligand. We compared the model training results achieved with the artificial pockets only, experimental pockets only, and a combination of both.

A subset of PDBbind complexes, which had been previously used in the analysis of DiffDock technique^8^, was used for model testing. We additionally removed all complexes containing the ligands with more than 100 heavy atoms in order to concentrate on the drug-like molecules.

### Model performance metrics

In a manner consistent with the EquiBind ^4^, TANKBind ^3^, and DiffDock ^8^ we used 25th, 50th, and 75th percentiles of symmetry-corrected^27^ Root Mean Square Deviation (RMSD) between expected and predicted ligand pose, together with the percentage of predictions below 5Å or 2Å RMSD. The centroid distances between the expected and predicted positions of the ligands were tracked using the same metrics.

The following additional metrics of the non-bond interactions were used:

- The fraction of favorable contacts (*F_fav_*) - total number of favorable interactions normalised by the number of ligand heavy atoms. The contribution of the hydrophobic interactions was accounted for with a weight of 1/6 due to their high relative abundance.
- The fraction of atoms involved in favorable contacts (*F_fav-atoms_*) - a fraction of the heavy ligand atoms participating in at least one non-bond interaction;
- The fraction of unfavorable contacts (*F_unfav_*) - total number of unfavorable interactions normalised by the number of ligand heavy atoms;

In order to evaluate the quality of predicted ligand poses, we determined the frequency of steric clashes between the ligand and protein atoms. The clash was defined as the distance between the atoms smaller than 70% of the sum of their respective Van der Waals radii.

### Model training and inference

The core architecture utilized in this study was the diffusion generative score model, which was adapted from DiffDock. This model is based on SE(3)-equivariant convolutional networks that operate on point clouds ^28,29^. The model utilizes pocket and ligand graph representations and takes into account the spatial arrangement of the atoms. It produces SE(3)-equivariant vectors that describe the ligand’s translations and rotations, along with SE(3)-invariant scalars per each compound’s rotatable bond. The input position of a ligand is subsequently altered based on the model’s output, resulting in the final conformation of the molecule within a binding site. During the training phase, the position of each input ligand in a complex undergoes modifications caused by translational, rotational, and torsional noise, which can be referred to as forward diffusion. The model then learns how to reverse the diffusion process. This method enables the generation of numerous alternative ligand poses during the inference process ^8^.

We adjust the DiffDock model inputs and architecture as follows:

- The atomic-level pocket graph is used instead of the residue-level graph of the whole protein.
- The categorical feature space of the ligand nodes was decreased significantly by narrowing down the number of atom types, number of neighbor heavy atoms, and atomic charges to those expected in the typical screening databases of small molecules.

The learning rate was set to 0.0125 based on initial tuning. During each epoch, the model is trained for 10,000 iterations with a batch size of 4. In our experimental conditions, the batch always consists of identical pocket and ligand components, whereas the level of diffusion noise varies between individual samples.

During the training based on artificial data, 10,000 ligands are randomly sampled from the preprocessed ZINC dataset at each epoch. After that, the artificial pockets are created for each ligand. Each epoch of training on experimental protein-ligand complexes is performed with 10,000 randomly sampled PDBbind entries. The training on a combined dataset was performed with a 4:1 ratio of artificial and experimental data. The models were trained for 25 epochs. The final production model was trained for 80 epochs using a combined dataset.

The model training, which is a GPU-intensive task, and pocket generation, which is a CPU-intensive task, were separated into distinct parallel workflows.

Given the generative nature of the model, it is possible to produce an infinite number of alternate ligand poses. The real-world model performance is thus sensitive to the scoring function, which is used for the pose ranking and selection. The developers of DiffDock employed a confidence model that takes into account all protein atoms and produces a confidence score of the ligand pose. In contrast, we employed a non-covalent interaction score in the binding pocket, which reduces the inference cost significantly. Our scoring function *S* is computed as follows:

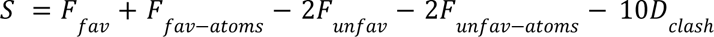

where *F_unfav-atoms_* is a fraction of the ligand heavy atoms participating in at least one unfavorable interaction, *D_clash_* is a sum of all distances, which are below the steric clashes threshold. The coefficients of the scoring function were adjusted empirically by visual inspection of the predicted protein-ligand complexes.

## Results and discussion

### Statistics of non-bond interactions in experimental and artificial protein-ligand complexes

Analysis of the high-quality PDBbind protein-ligand dataset revealed a total of 343,784 intermolecular non-covalent interactions. As anticipated, the hydrophobic contacts were the predominant kind of interaction, with a total of 267,774 atom pairs observed. The overwhelming majority of these interactions occurred between hydrophobic carbon or sulfur atoms. The hydrogen bonds were the second most prevalent form of interaction, occurring around once for every five hydrophobic pairs. The remaining contacts constituted less than 3% of the overall count (Figure 2, Table S2).

**Figure 2.**
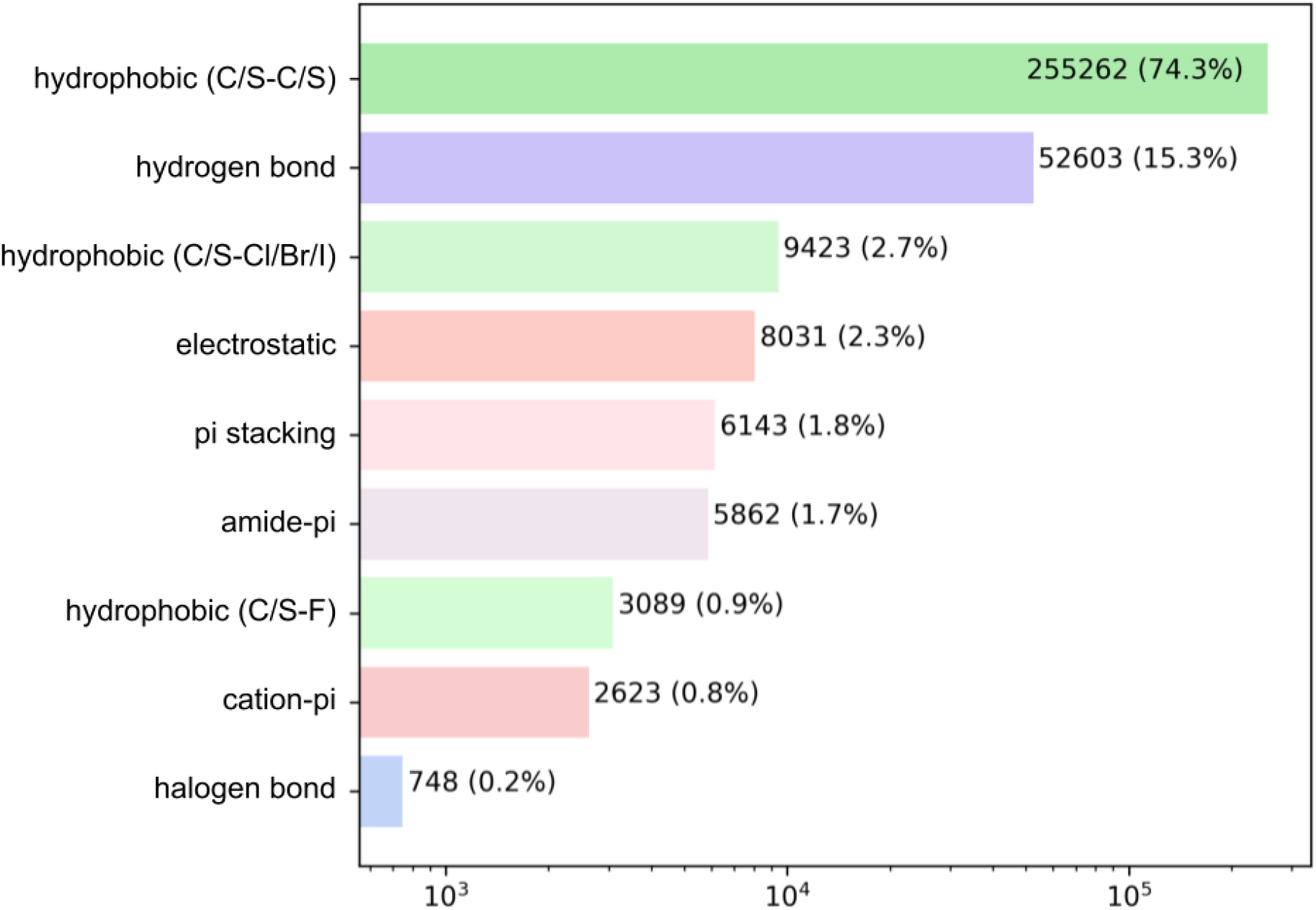
Probability distribution of the non-bond protein-ligand interactions in the PDBbind complexes. Every bar is labeled with the total number of interactions and its fraction. Note the log scale of the X-axis.

The total number of non-bond interaction entries in the artificial pockets involving the same ligands amounted to 453,593. We compared the distributions of the parameters for each type of interaction between real and artificial pockets. For each interaction type, we also computed the probability of the involvement of particular amino acids in the binding pocket.

Figure 2 shows the statistics of hydrophobic interactions in real and artificial binding pockets

There is a reasonably good correspondence between the distributions, but the distributions of distances are systematically smoother and more monotonous for artificial pockets in comparison to real ones. This is an artifact caused by the frequent formation of “unintended” hydrophobic contacts while generating other types of interactions due to the abundance of hydrophobe atoms in both amino acids and small molecules. The distributions of interactions among amino acids are very similar except the notably increased involvement of PHE and TRP in artificial pockets. This is explained by the overlap between hydrophobic and pi stacking interactions, which is not checked in the algorithm for the sake of computational efficiency.

Figure 3 shows the statistics of pi stacking interactions. There are two types of these interactions: parallel (true pi stacking) and T-shaped (aromatic-pi interactions). The distance distributions of both types of interactions are remarkably similar in real and artificial pockets as well as the involvements of different aromatic amino acids. However, the theta angle distributions of parallel interactions for artificial pockets are shifted toward 90° by 10-15 degrees in comparison to experimental ones since the generation algorithm only checks if the aromatic ring has an angle within a given range.

**Figure 3.**
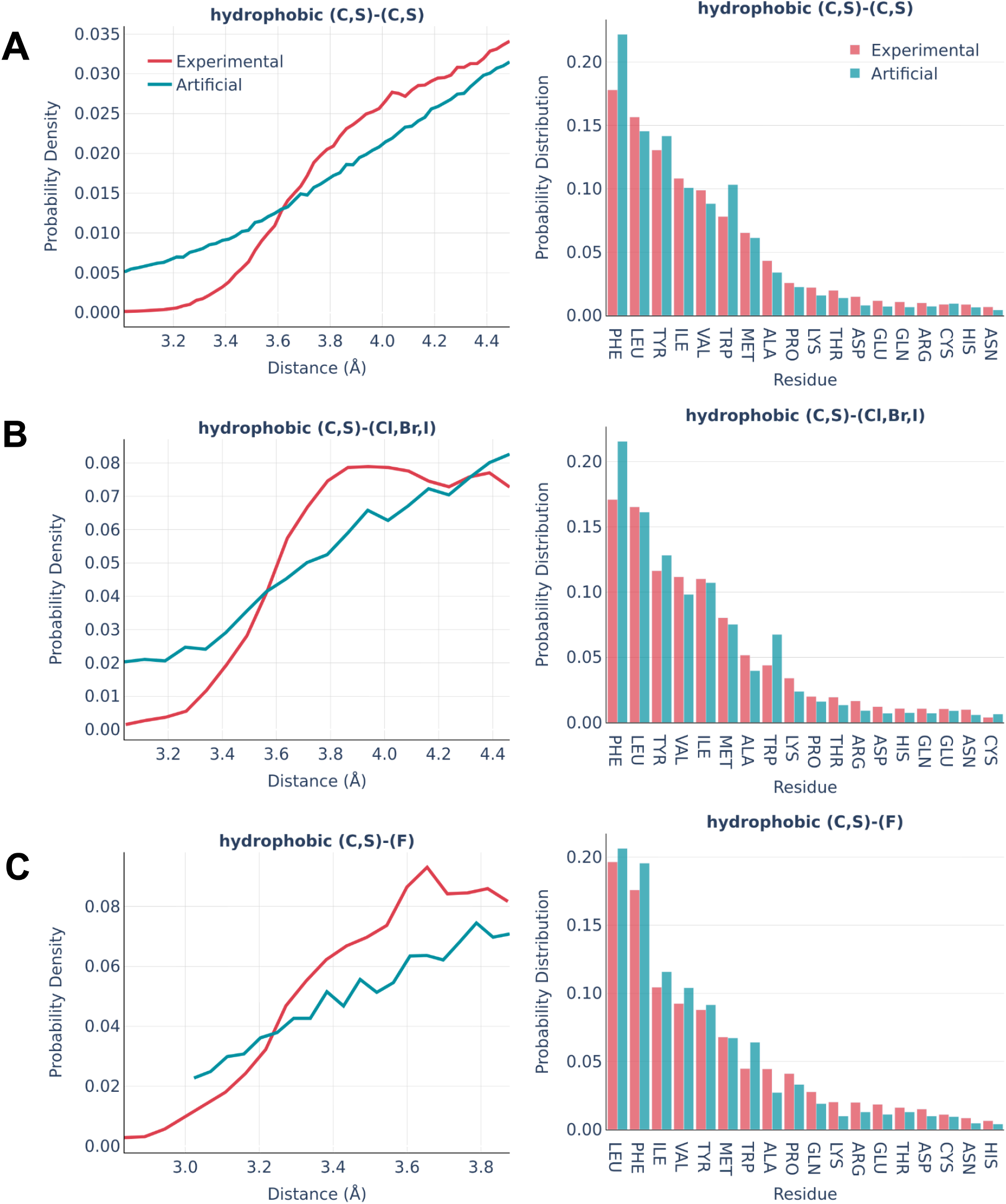
Statistics of hydrophobic interactions in the PDBbind and artificial pockets involving (A) C or S atoms of the protein a and the ligand; (B) Br, Cl or I atoms of the ligand; (C) F atoms of the ligand. The distance distributions are shown in the left columns and the pocket residues occurrence ratio is shown in the right column.

Figure 4 shows the statistics of amide-pi interactions, which could also be classified into parallel and T-shaped.

**Figure 3.**
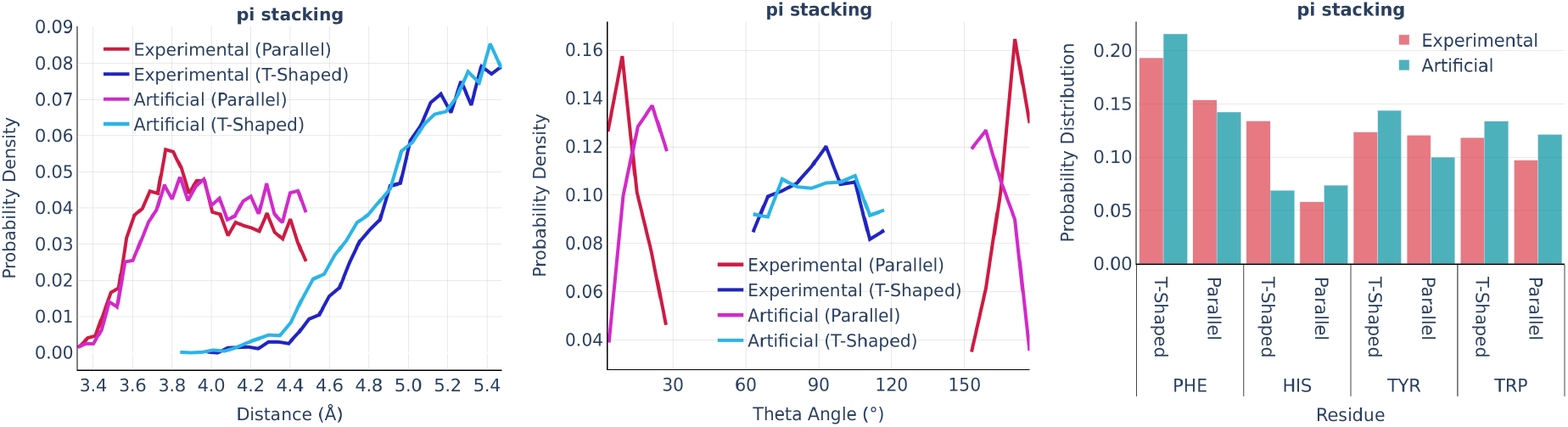
pi stacking: the distance distribution (left), Theta angle distribution (middle), and pocket residues frequency (right) in the PDBbind and artificial pockets.

**Figure 4.**
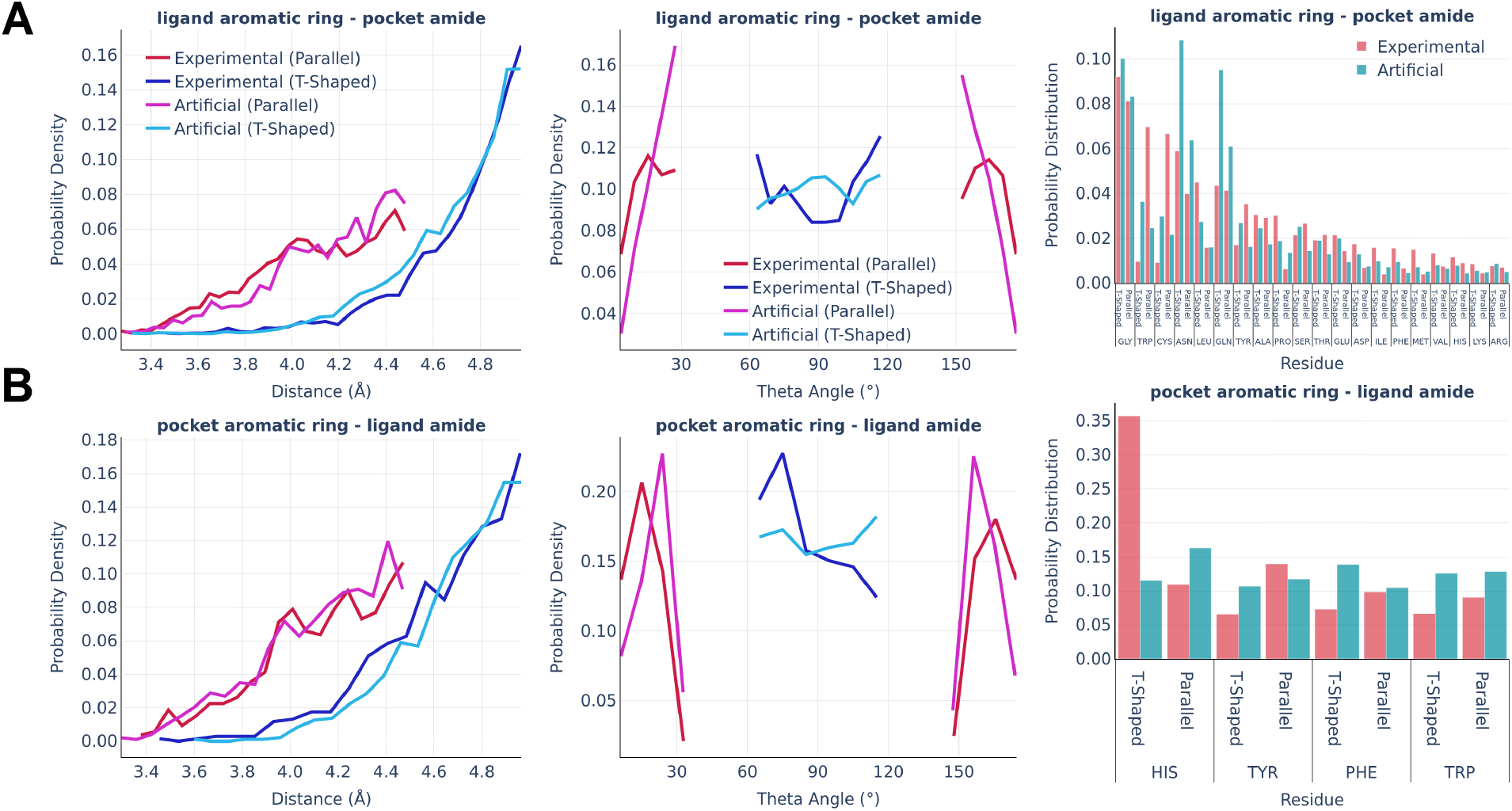
Amide-pi interactions: the distance distribution (left), Theta angle distribution (middle), and pocket residues frequency (right) in the PDBbind and artificial pockets for the ligand aromatic ring - pocket amide contacts (A) and pocket aromatic ring - ligand amide contacts (B).

Similarly to pi stacking interactions, the distance distributions are very similar between real and artificial pockets, while the theta angles in artificial pockets are somewhat shifted towards 90°. The primary residues serving as donors of the amide group were found to be glycine, which exposes the peptide bond due to the lack of a side chain, as well as asparagine and glutamine, which possess an amide group at the terminus of their side chains. The predominant residue bearing an aromatic ring in the amide-pi linkages is HIS, which accounts for the majority of experimentally observed T-shaped interactions. There are significantly less HIS contacts in artificial pockets because unfavorable donor-donor contacts between the nitrogens of the HIS ring and amide group were omitted during the pocket generation.

The distributions of the hydrogen bonds are shown in Figure 6.

**Figure 6.**
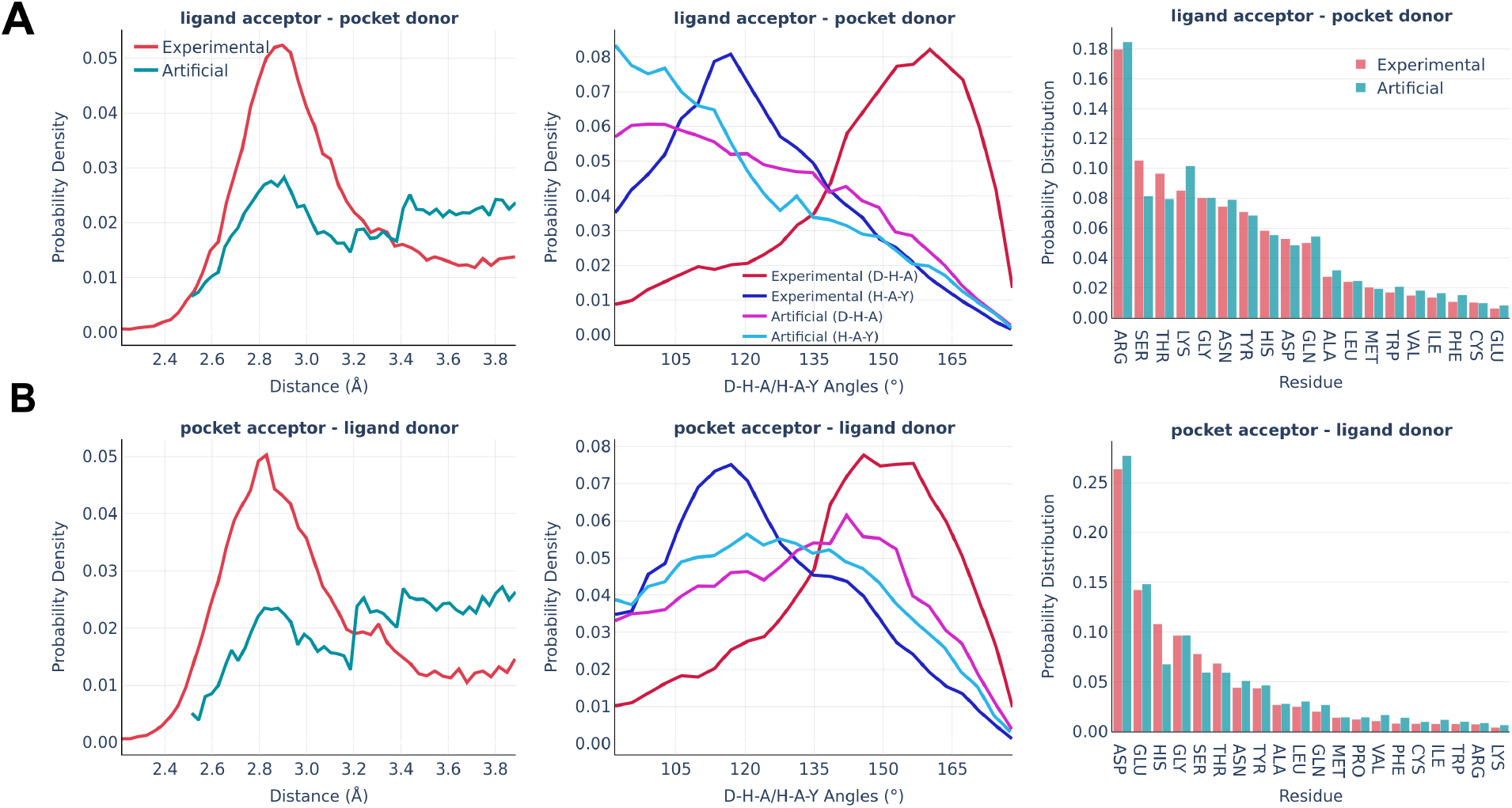
Hydrogen bonds: the distance distribution (left), D-H-A/H-A-Y angles distribution (middle), and pocket residues frequency (right) in the PDBbind and artificial pockets for the ligand acceptor - pocket donor contacts (A) and pocket acceptor - ligand donor contacts (B). In the D-H-A/H-A-Y angles, D is the donor atom covalently bound to the hydrogen; H is a hydrogen atom; A is an acceptor atom; Y is a heavy covalently bound neighbor of the acceptor atom.

Hydrogen bonding imposes the biggest challenge in terms of artificial pocket generation. It is clearly seen there are the biggest discrepancies between real and artificial pockets in terms of the hydrogen bonds’ parameters, which are especially visible for the angle distributions. In general, our algorithm tends to underestimate the D-H-A angles (the significance of the angle distribution was deliberately reduced to speed up the algorithm) and doesn’t capture the distance peak at 2.8-2.9 Å, which is observed in real pockets, while generating significantly more long h-bonds (short h-bonds tend to be discarded more often during the generation as they often lead to steric clashes). These compromises allow us to keep the algorithm fast enough for routine practical usage. At the same time, the involvement of different amino acids is reproduced remarkably well in artificial pockets.

The halogen bonds are the least frequent type of interaction in real binding pockets. Their distributions are shown in Figure 7. Taking into account the small number of observed interactions of this kind, the experimental and artificial distributions are sufficiently similar to each other.

**Figure 7.**
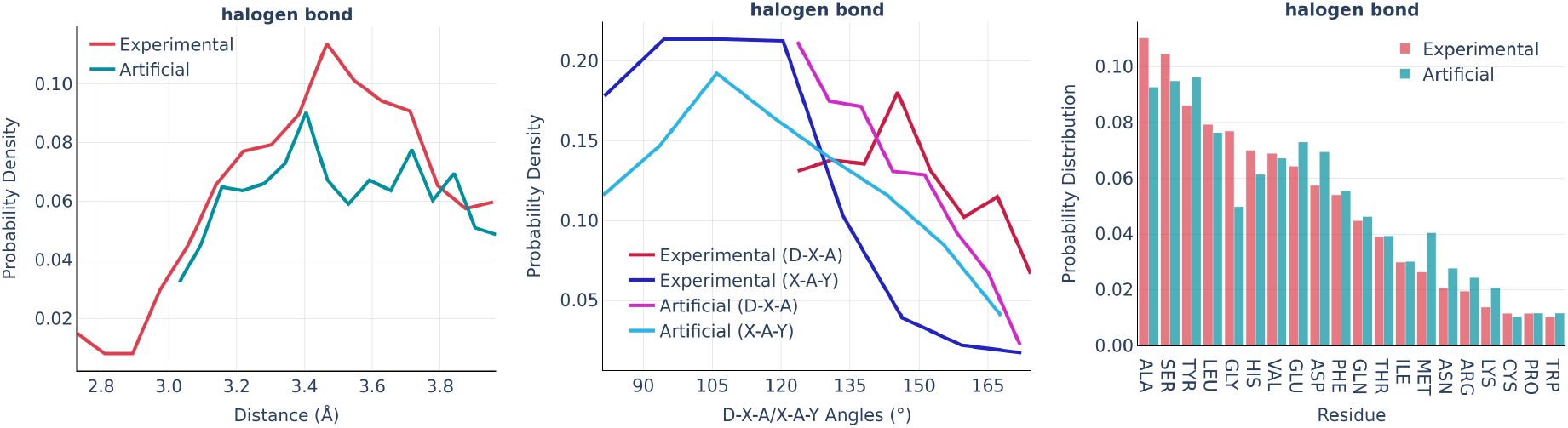
Halogen bonds: the distance distribution (left), D-X-A/X-A-Y angles distribution (middle), and pocket residue frequencies (right) in the PDBbind and artificial pockets. In the D-X-A/X-A-Y angles, D is the donor atom covalently bound to the halogen; X is a halogen atom; A is the halogen bond acceptor atom; Y is a heavy covalently bound neighbor of the acceptor atom.

The electrostatic interactions were analyzed separately for the “ligand anion - pocket cation” and “pocket anion - ligand cation” pairs (Figure 8).

**Figure 8.**
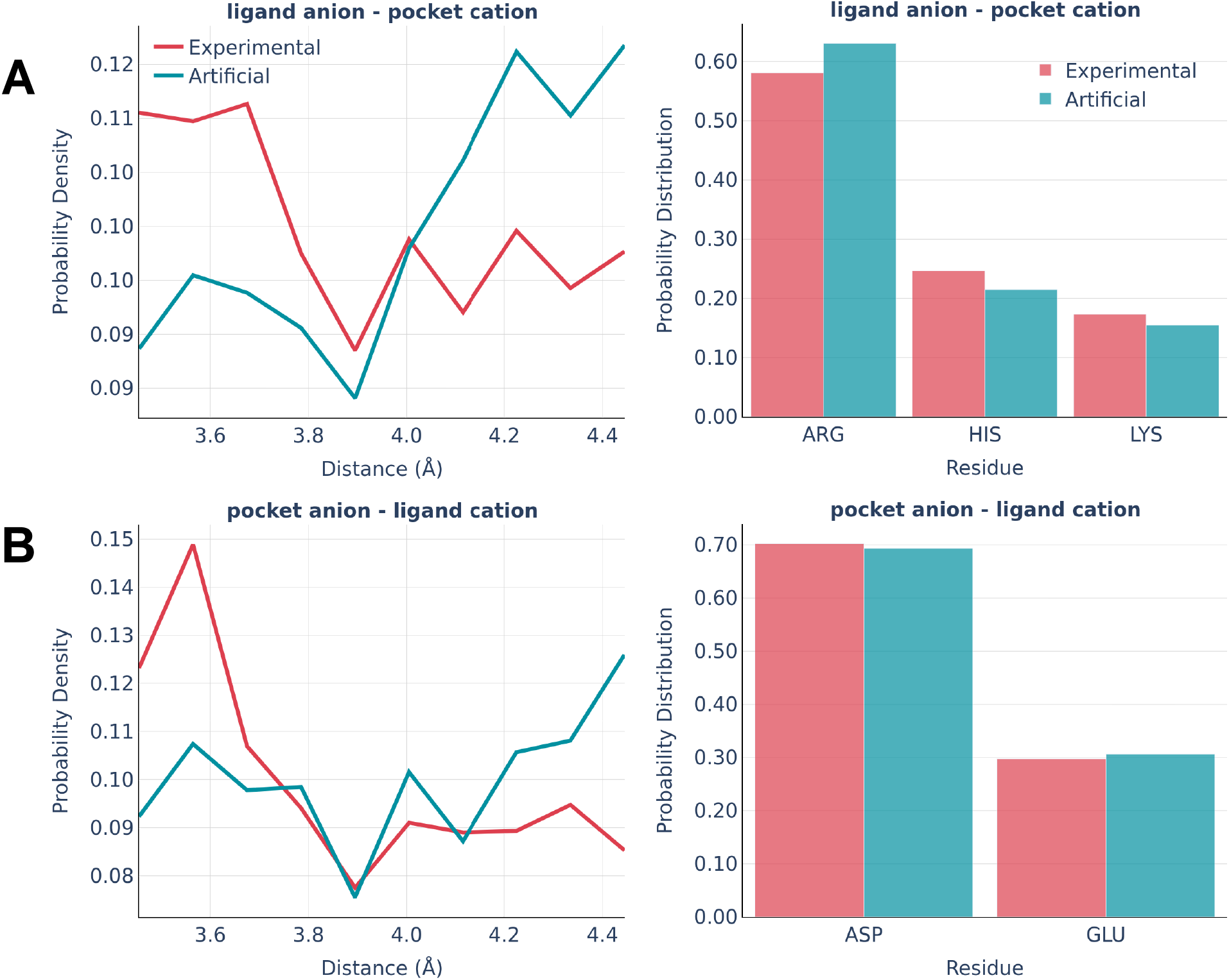
Electrostatic interactions: the distance distribution (left) and pocket residues frequency (right) in the PDBbind and artificial pockets for the ligand anion - pocket cation (A) and pocket anion - ligand cation (B).

The experimentally observed involvement of charged amino acids is reproduced almost ideally in artificial pockets, while the distance distributions are somewhat different in real and artificial pockets. Artificial pockets possess more iterating pairs at larger distances than real ones, which could be explained by the increased complexity of fitting pocket residues in the close proximity of a small molecule. Such placement often leads to steric clashes and thus is often discarded by the algorithm.

The last type of interaction is cation-pi pairs (Fig. 9). They are rather minor and the distributions of their parameters are very similar in real and artificial pockets without any significant feature worth commenting on.

**Figure 9.**
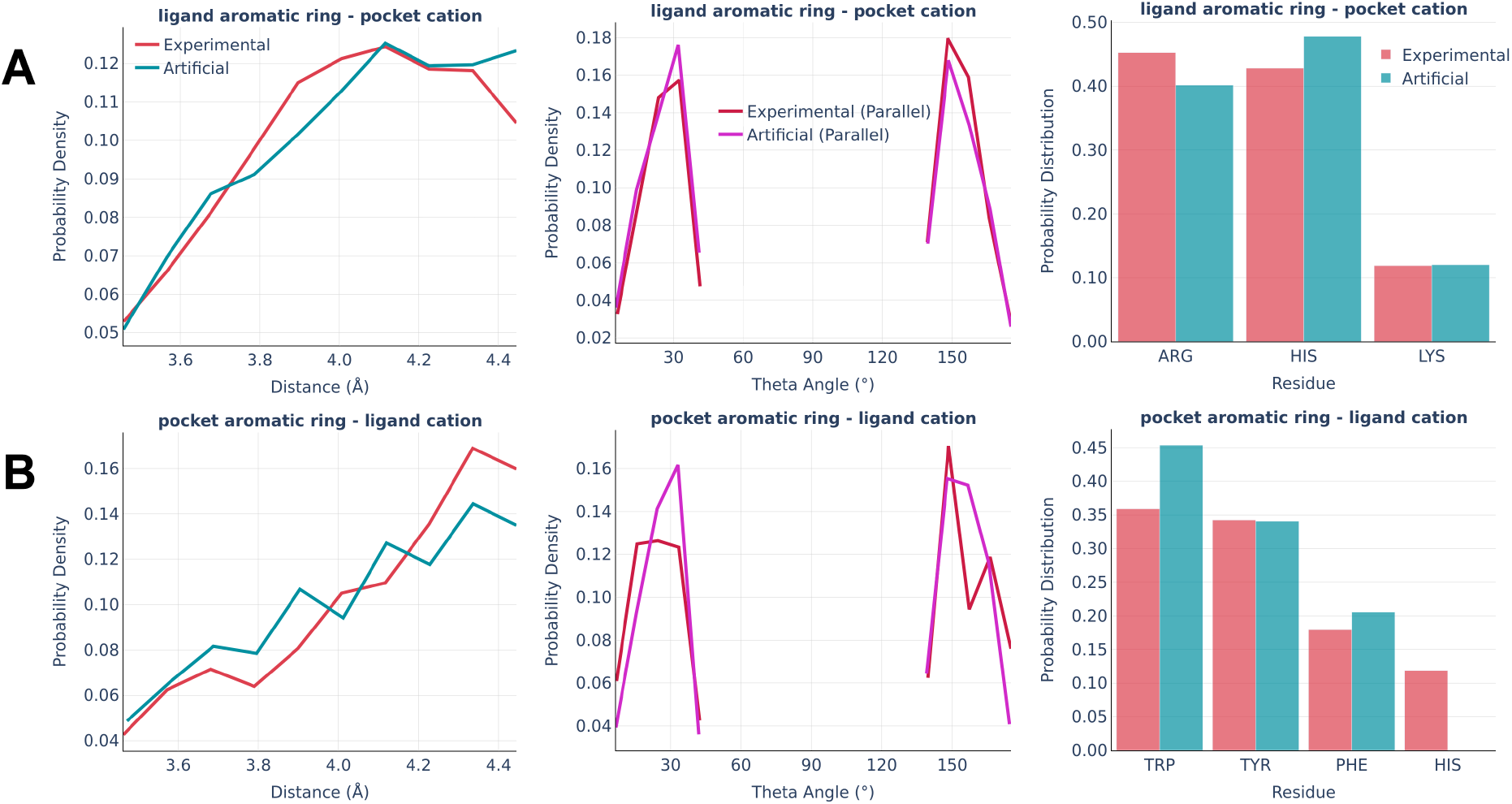
Cation-pi interactions: the distance distribution (left), Theta angle distribution (middle), and pocket residues frequency (right) in the PDBbind and artificial pockets for the ligand aromatic ring - pocket cation (A) and pocket aromatic ring - ligand cation (B).

### Impact of artificial data on model performance

To evaluate the potential influence of data augmentation using artificial protein-ligand complexes on model performance, we compared three models. The first model was exclusively trained on experimental protein-ligand complexes, the second model was trained on the artificially generated pockets with corresponding small molecules, and the third model was trained on the combination of both. The metrics are reported for the top-1 predicted complex and the best of the top-5 predicted complexes based on custom scoring function S (see the Methods section).

Table 2 illustrates that the inclusion of artificial or combined data in the training process led to enhancements in both the RMSD and Centroid Distance metrics, as compared to the model trained exclusively on experimental complexes. Unexpectedly, the training of the artificial dataset is superior to the combined dataset in certain metrics, particularly when the generation of 40 poses for each pocket was used.

**Table 2.** The symmetry-corrected Root Mean Square Deviation (RMSD) and Centroid Distance metrics between predicted and real ligand positions in the PDBbind test complexes. The models generated either 10 or 40 ligand poses. The best of top 5 poses is picked based on the lowest RMSD. ↑ - means the higher the better; ↓ - means the lower the better.

| Dataset | RMSD |  |  |  |  | Centroid Distance |  |  |  |  |
| --- | --- | --- | --- | --- | --- | --- | --- | --- | --- | --- |
|  | Percentiles ↓ |  |  | % below threshold ↑ |  | Percentiles ↓ |  |  | % below threshold ↑ |  |
|  | 25th | 50th | 75th | 5Å | 2Å | 25th | 50th | 75th | 5Å | 2Å |

| 10 Samples, Top 1 |  |  |  |  |  |  |  |  |  |  |
| --- | --- | --- | --- | --- | --- | --- | --- | --- | --- | --- |
| Experimental | 4.08 | 5.51 | 7.45 | 40.93 | 5.02 | 1.02 | 1.75 | 2.7 | 94.59 | 57.92 |
| Artificial | **3.69** | 5.75 | 7.39 | 40.15 | 5.02 | **1** | **1.46** | **2.19** | 97.68 | **69.88** |
| Combined | 3.76 | **5.13** | **7.12** | **45.95** | **7.34** | 1.05 | 1.69 | 2.33 | **98.07** | 63.32 |
| 10 Samples, Top 5 |  |  |  |  |  |  |  |  |  |  |
| Experimental | 2.68 | 3.72 | 4.64 | 82.24 | 11.2 | 0.95 | 1.37 | 1.97 | **99.61** | 75.29 |
| Artificial | 2.49 | 3.47 | 4.6 | 81.08 | 16.99 | **0.78** | **1.24** | **1.86** | 98.84 | **80.31** |
| Combined | **2.4** | **3.31** | **4.55** | **85.33** | **17.37** | 0.81 | 1.29 | 1.88 | 99.23 | 77.99 |
| 40 Samples, Top 1 |  |  |  |  |  |  |  |  |  |  |
| Experimental | 3.89 | 5.54 | 7.36 | 40.93 | 4.63 | 1.01 | 1.73 | 2.48 | 96.14 | 57.92 |
| Artificial | **3.47** | 5.46 | 7.25 | **47.1** | **7.72** | **1** | **1.39** | **2.09** | **97.68** | **72.59** |
| Combined | 3.86 | **5.2** | **7.16** | 45.95 | 6.18 | 1.05 | 1.57 | 2.42 | **97.68** | 64.09 |
| 40 Samples, Top 5 |  |  |  |  |  |  |  |  |  |  |
| Experimental | 2.37 | 3.35 | 4.6 | 82.24 | 15.06 | 0.85 | 1.19 | **1.69** | **99.61** | 81.08 |
| Artificial | **1.99** | **3.08** | **4.3** | **85.33** | **25.1** | **0.73** | **1.11** | **1.69** | **99.61** | **82.63** |
| Combined | 2.09 | 3.3 | 4.6 | 82.24 | 23.94 | 0.8 | 1.26 | 1.99 | 98.84 | 75.29 |

The combined dataset of protein-ligand complexes exhibited superior performance compared to the other two datasets in terms of both favorable and unfavorable non-covalent interactions, as illustrated in Table 3. Additionally, it was observed that the integration of the combined data for training purposes led to a decrease in the proportion of samples exhibiting steric clashes, as shown in Table 4.

**Table 3.** The Favorable Rate, Favorable Rate Uniq, and Unfavorable Rate for the predicted PDBbind test complexes. The models generated either 10 or 40 samples. The best of the top 5 candidates are picked based on the lowest RMSD. ↑ - means the higher the better; ↓ - means the lower the better.

|  | Favorable Rate |  |  | Favorable Rate Uniq |  |  | Unfavorable Rate |  |  |
| --- | --- | --- | --- | --- | --- | --- | --- | --- | --- |
|  | Percentiles ↑ |  |  | Percentiles ↑ |  |  | Percentiles ↓ |  |  |
| Dataset | 25th | 50th | 75th | 25th | 50th | 75th | 25th | 50th | 75th |
| 10 Samples, Top 1 |  |  |  |  |  |  |  |  |  |
| Experimental | **0.27** | 0.37 | 0.51 | 0.38 | 0.53 | 0.67 | **0** | 0.08 | 0.17 |
| Artificial | **0.27** | **0.39** | 0.53 | 0.38 | 0.53 | 0.67 | **0** | **0.07** | **0.15** |
| Combined | **0.27** | 0.38 | **0.54** | **0.41** | **0.54** | **0.68** | **0** | **0.07** | **0.15** |
| 10 Samples, Top 5 |  |  |  |  |  |  |  |  |  |
| Experimental | 0.28 | 0.38 | 0.52 | 0.35 | 0.53 | 0.66 | 0.04 | 0.1 | 0.23 |
| Artificial | **0.3** | 0.39 | 0.52 | 0.38 | 0.52 | **0.68** | 0.04 | 0.1 | 0.21 |
| Combined | **0.3** | **0.4** | **0.56** | **0.39** | **0.55** | 0.67 | **0.03** | **0.09** | **0.18** |
| 40 Samples, Top 1 |  |  |  |  |  |  |  |  |  |
| Experimental | 0.29 | 0.37 | 0.54 | 0.4 | 0.54 | **0.71** | **0** | 0.06 | 0.14 |
| **Artificial** | 0.29 | 0.41 | 0.58 | **0.42** | **0.58** | **0.71** | **0** | **0.04** | **0.12** |
| **Combined** | **0.31** | **0.43** | **0.59** | **0.42** | **0.58** | **0.71** | **0** | 0.05 | **0.12** |
| **40 Samples, Top 5** |  |  |  |  |  |  |  |  |  |
| **Experimental** | 0.26 | 0.4 | 0.54 | 0.39 | 0.53 | 0.69 | 0.03 | 0.09 | 0.18 |
| **Artificial** | 0.29 | 0.41 | **0.55** | 0.39 | 0.54 | 0.7 | **0** | 0.07 | **0.14** |
| **Combined** | **0.32** | **0.43** | **0.55** | **0.41** | **0.56** | **0.71** | **0** | **0.06** | **0.14** |

**Table 4.** The percentage of samples within the predicted PDBbind test complexes exhibiting at least one steric clash. The models generated 40 ligand poses. The best of top-5 candidates are picked based on the lowest number of clashes.

| **Dataset** | **Steric Clashes** |  |
| --- | --- | --- |
|  | **Top 1** | **Top 5** |
| **Experimental** | 33.59% | 30.12% |
| **Artificial** | 30.50% | 26.64% |
| **Combined** | **28.19%** | **23.94%** |

A positive correlation was identified between the amount of small molecule heavy atoms and the metrics such as RMSD and Steric Clashes. Additionally, a negative correlation was found between the count of atoms and favorable non-bond interaction rates, as depicted in Figure S2. This indicates that the model performance decreases with the increase of ligand size.

The final production PocketCFDM model was trained for 80 epochs using high-quality experimental data (used for the non-bond interactions statistics retrieval) and artificial samples, similar to the aforementioned combined dataset settings. A significant reduction in the occurrence of protein-ligand steric clashes was observed, with percentages of 19.31% and 13.90% for the top-1 and best of top-5 samples, respectively. This is a nearly 10% decrease compared to the most optimal model trained for 25 epochs (Table 4). There has been no substantial improvement in other metrics.

### Comparison with DiffDock

We also performed a detailed comparison between PocketCFDM and DiffDock, which is currently considered as the most accurate AI technique for predicting ligand binding poses (Table 5).

**Table 5.**
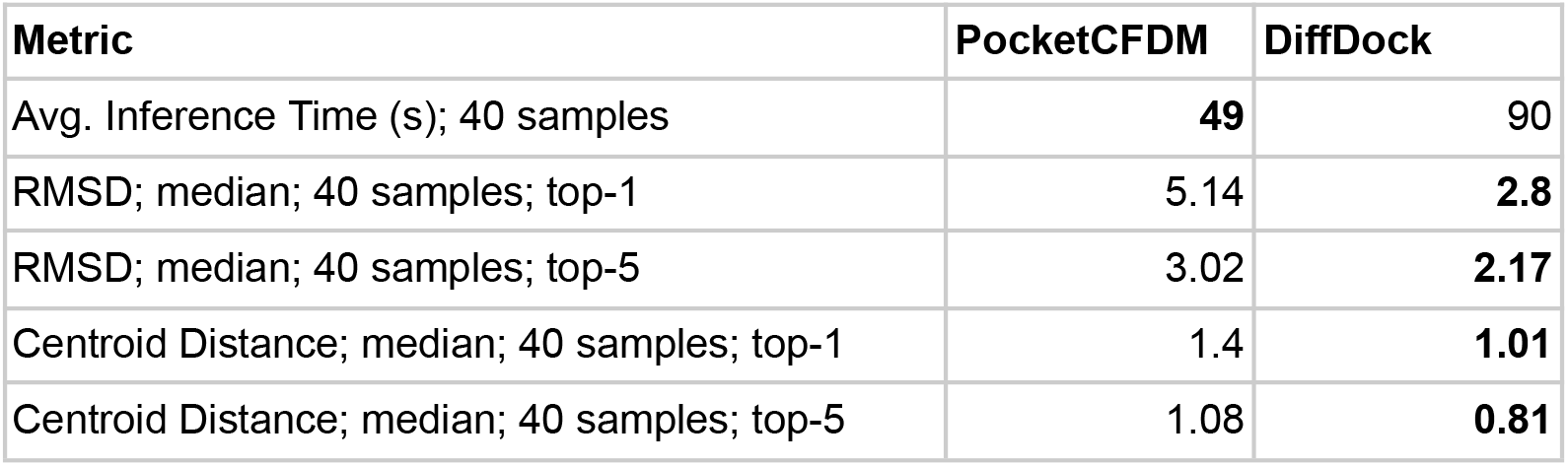
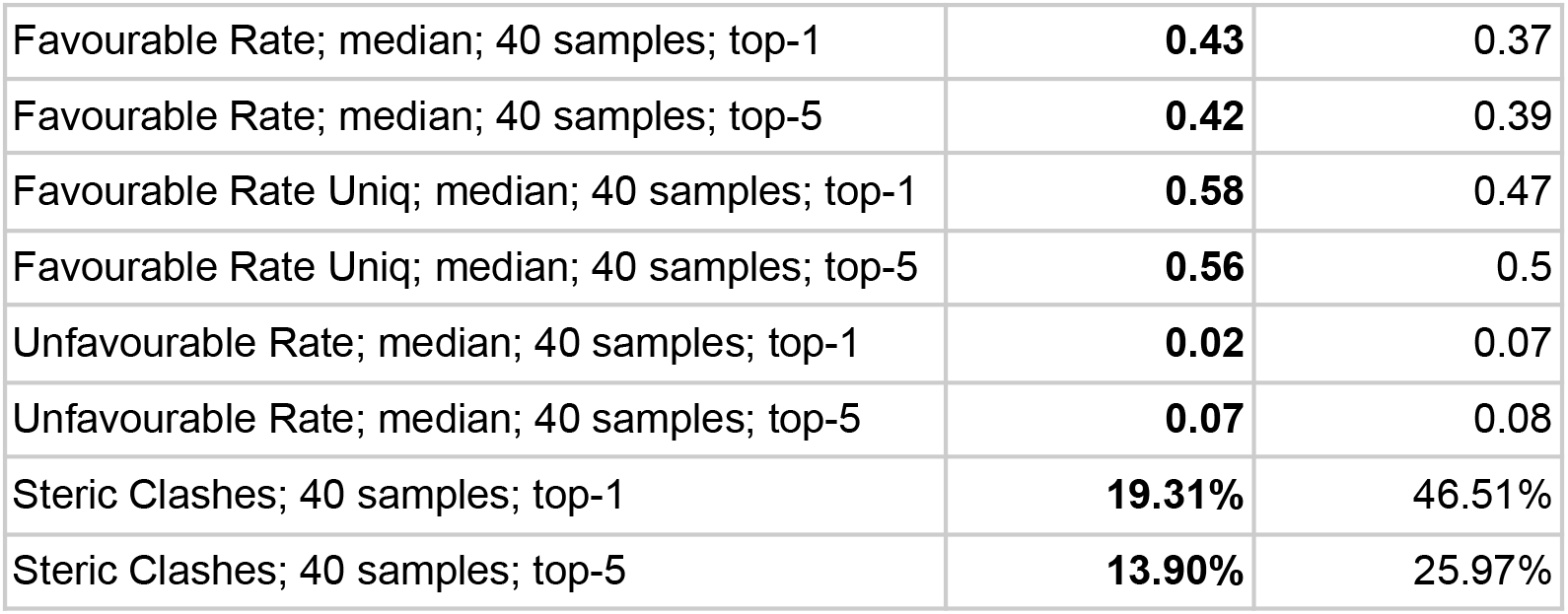
Comparison between the DiffDock and PocketCFDM on the test PDBbind dataset.

DiffDock exhibited superior accuracy in terms of RMSD and Centroid Distance. In contrast, PocketCFDM exhibited superior outcomes in terms of favorable and unfavorable non-covalent interactions, as well as a notably lower incidence of steric clashes. It was expected due to the divergence in the scoring algorithms employed to evaluate and rank the generated samples. The DiffDock approach utilized a confidence model that was trained to prioritize samples with lower RMSD to the actual ligand pose. However, in our context, the scoring approach was more focused on the number of non-covalent contacts and the absence of steric clashes, without considering the specific conformation of the ligand pose. Another notable difference pertaining to the disparity in scoring functions is the diversity observed within anticipated poses. The median RMSD values between the alternative top-5 samples in the PocketCFDM and DiffDock methods were 2.59 and 1.29 respectively. Additionally, it was noted that the mean inference time for the PocketCFDM was approximately 1.8 times quicker. This is attributed mostly to the reduced size of the protein graph and the decrease in the graph’s node feature space.

### Limitations and perspectives

The primary area of enhancement of the PocketCFDM model lies in inference time, which is currently still too large for efficient practical deployment. The mean time required to predict 40 ligand poses is around ∼50 seconds at 24 GB NVIDIA L4 GPU. The inference time could be improved significantly by replacing computationally intensive EGNN blocks with more efficient alternatives like GVP. The inclusion of the artificial pockets into the training of even simpler regression-based techniques, such as EquiBind and TANKBind, could also be beneficial since these architectures are generally more sensitive to the number of distinct training samples. It is possible to explore the usage of multiple augmentations of the same input instead of directly including specific symmetries and equivariances into Graph Neural Network (GNN) designs. The utilization of such augmentation is popular in Convolutional Neural Networks (CNNs). Moreover, it demonstrated promising outcomes in the domain of geometric graph learning^30^. This strategy may potentially increase the model training time while resulting in reduced inference time.

Another notable drawback of the present proof-of-the-principle implementation is the possibility of ligand self-intersections (intramolecular clashes), which have to be filtered out during the post-processing steps. Incorporating intramolecular and intermolecular clashes into the loss function during the training process could potentially address this problem.

Also, we believe that incorporating larger ligands into the training process will address the challenges encountered with relatively large compounds. The ZINC20 dataset, which was utilized in this work, has only 2.5% of molecules larger than 40 heavy atoms, which makes them underrepresented during the model training. Although the median size of the ligands in this dataset is 25 heavy atoms, which is quite common for the datasets of drug-like molecules, a higher percentage of large ligands may enhance the inference capabilities for larger compounds while maintaining the same level of performance for smaller molecules.

## Conclusions

In this work, we introduced the PocketCFDM generative diffusion model for predicting the poses of small molecules in the protein binding pockets. The model is trained using an innovative approach of data augmentation, which involves the construction of a large number of artificial binding pockets that follow the same statistical patterns of non-bond interactions as the real ones. In order to construct such artificial pockets we thoroughly evaluated the statistical characteristics of non-bond interactions in the real protein-ligand complexes and designed an algorithmic approach that reproduces them in artificial pockets, which are built around given small molecule conformers. The performance of the models trained on experimental data only, artificial data only, and a combination of both was evaluated and compared to the current state-of-the-art ML model for binding pose prediction DiffDock. It is shown that the inclusion of artificial binding pockets into the model training resulted in significant increase of model performance. Particularly, PocketCFDM outperforms DiffDock in terms of the quality of non-bond interactions, the number of steric clashes, and the inference speed. The perspectives of further improvements of PocketCFDM are discussed. The inference code and final model weights are publicly available in the GitHub repository (https://github.com/vtarasv/pocket-cfdm.git).

## References

1. Li, X., Li, Y., Cheng, T., Liu, Z. & Wang, R. Evaluation of the performance of four molecular docking programs on a diverse set of protein-ligand complexes. J. Comput. Chem. 31, 2109–2125 (2010).

2. Ghasemi, J. B., Abdolmaleki, A. & Shiri, F. Molecular Docking Challenges and Limitations. in Pharmaceutical Sciences: Breakthroughs in Research and Practice 770–794 (IGI Global, 2017). doi:10.4018/978-1-5225-1762-7.ch030.

3. Lu, W., et al. TANKBind: Trigonometry-Aware Neural NetworKs for Drug-Protein Binding Structure Prediction. http://biorxiv.org/lookup/doi/10.1101/2022.06.06.495043 (2022) doi:10.1101/2022.06.06.495043.

4. Stärk, H., Ganea, O.-E., Pattanaik, L., Barzilay, R. & Jaakkola, T. EquiBind: Geometric Deep Learning for Drug Binding Structure Prediction. Preprint at 10.48550/arXiv.2202.05146 (2022).

5. Jing, B., Eismann, S., Suriana, P., Townshend, R. J. L. & Dror, R. Learning from Protein Structure with Geometric Vector Perceptrons. (2020) doi:10.48550/ARXIV.2009.01411.

6. Jing, B., Eismann, S., Soni, P. N. & Dror, R. O. Equivariant Graph Neural Networks for 3D Macromolecular Structure. (2021) doi:10.48550/ARXIV.2106.03843.

7. Ganea, O.-E. et al. Independent SE(3)-Equivariant Models for End-to-End Rigid Protein Docking. (2021) doi:10.48550/ARXIV.2111.07786.

8. Corso, G., Stärk, H., Jing, B., Barzilay, R. & Jaakkola, T. DiffDock: Diffusion Steps, Twists, and Turns for Molecular Docking. (2022) doi:10.48550/ARXIV.2210.01776.

9. Wang, R., Fang, X., Lu, Y. & Wang, S. The PDBbind Database: Collection of Binding Affinities for Protein−Ligand Complexes with Known Three-Dimensional Structures. J. Med. Chem. 47, 2977–2980 (2004).

10. Liu, Z. et al. PDB-wide collection of binding data: current status of the PDBbind database. Bioinformatics 31, 405–412 (2015).

11. Petrovski, Ž. H., Hribar-Lee, B. & Bosnić, Z. CAT-Site: Predicting Protein Binding Sites Using a Convolutional Neural Network. Pharmaceutics 15, 119 (2022).

12. Kandel, J., Tayara, H. & Chong, K. T. PUResNet: prediction of protein-ligand binding sites using deep residual neural network. J. Cheminformatics 13, 65 (2021).

13. Volkov, M. et al. On the Frustration to Predict Binding Affinities from Protein–Ligand Structures with Deep Neural Networks. J. Med. Chem. 65, 7946–7958 (2022).

14. The UniProt Consortium. UniProt: a worldwide hub of protein knowledge. Nucleic Acids Res. 47, D506–D515 (2019).

15. Reymond, J.-L., Van Deursen, R., Blum, L. C. & Ruddigkeit, L. Chemical space as a source for new drugs. MedChemComm 1, 30 (2010).

16. Bissantz, C., Kuhn, B. & Stahl, M. A Medicinal Chemist’s Guide to Molecular Interactions. J. Med. Chem. 53, 5061–5084 (2010).

17. Ferreira De Freitas, R. & Schapira, M. A systematic analysis of atomic protein–ligand interactions in the PDB. MedChemComm 8, 1970–1981 (2017).

18. Wilcken, R., Zimmermann, M. O., Lange, A., Joerger, A. C. & Boeckler, F. M. Principles and Applications of Halogen Bonding in Medicinal Chemistry and Chemical Biology. J. Med. Chem. 56, 1363–1388 (2013).

19. Kuhn, B., Gilberg, E., Taylor, R., Cole, J. & Korb, O. How Significant Are Unusual Protein–Ligand Interactions? Insights from Database Mining. J. Med. Chem. 62, 10441–10455 (2019).

20. Wójcikowski, M., Zielenkiewicz, P. & Siedlecki, P. Open Drug Discovery Toolkit (ODDT): a new open-source player in the drug discovery field. J. Cheminformatics 7, 26 (2015).

21. Jubb, H. C. et al. Arpeggio: A Web Server for Calculating and Visualising Interatomic Interactions in Protein Structures. J. Mol. Biol. 429, 365–371 (2017).

22. Adasme, M. F. et al. PLIP 2021: expanding the scope of the protein–ligand interaction profiler to DNA and RNA. Nucleic Acids Res. 49, W530–W534 (2021).

23. Tien, M. Z., Sydykova, D. K., Meyer, A. G. & Wilke, C. O. PeptideBuilder: A simple Python library to generate model peptides. PeerJ 1, e80 (2013).

24. Irwin, J. J. et al. ZINC20—A Free Ultralarge-Scale Chemical Database for Ligand Discovery. J. Chem. Inf. Model. 60, 6065–6073 (2020).

25. Sterling, T. & Irwin, J. J. ZINC 15 – Ligand Discovery for Everyone. J. Chem. Inf. Model. 55, 2324–2337 (2015).

26. Bento, A. P. et al. An open source chemical structure curation pipeline using RDKit. J. Cheminformatics 12, 51 (2020).

27. Meli, R. & Biggin, P. C. spyrmsd: symmetry-corrected RMSD calculations in Python. J. Cheminformatics 12, 49 (2020).

28. Thomas, N., et al. Tensor field networks: Rotation- and translation-equivariant neural networks for 3D point clouds. (2018) doi:10.48550/ARXIV.1802.08219.

29. Geiger, M. & Smidt, T. e3nn: Euclidean Neural Networks. (2022) doi:10.48550/ARXIV.2207.09453.

30. FAENet: Frame Averaging Equivariant GNN for Materials Modeling | OpenReview. https://openreview.net/forum?id=HRDRZNxQXc.

