## Supplementary Information for "Boosting performance of generative diffusion model for molecular docking by training on artificial binding pockets"

Supporting Information

Taras Voitsitskyi, Volodymyr Bdzhola, Roman Stratiichuk, Ihor Koleiev, Zakhar Ostrovsky, Volodymyr Vozniak, Ivan Khropachov, Pavlo Henitsoi, Leonid Popryho, Roman Zhytar, Semen Yesylevskyy, Alan Nafiiev, Serhii Starosyla

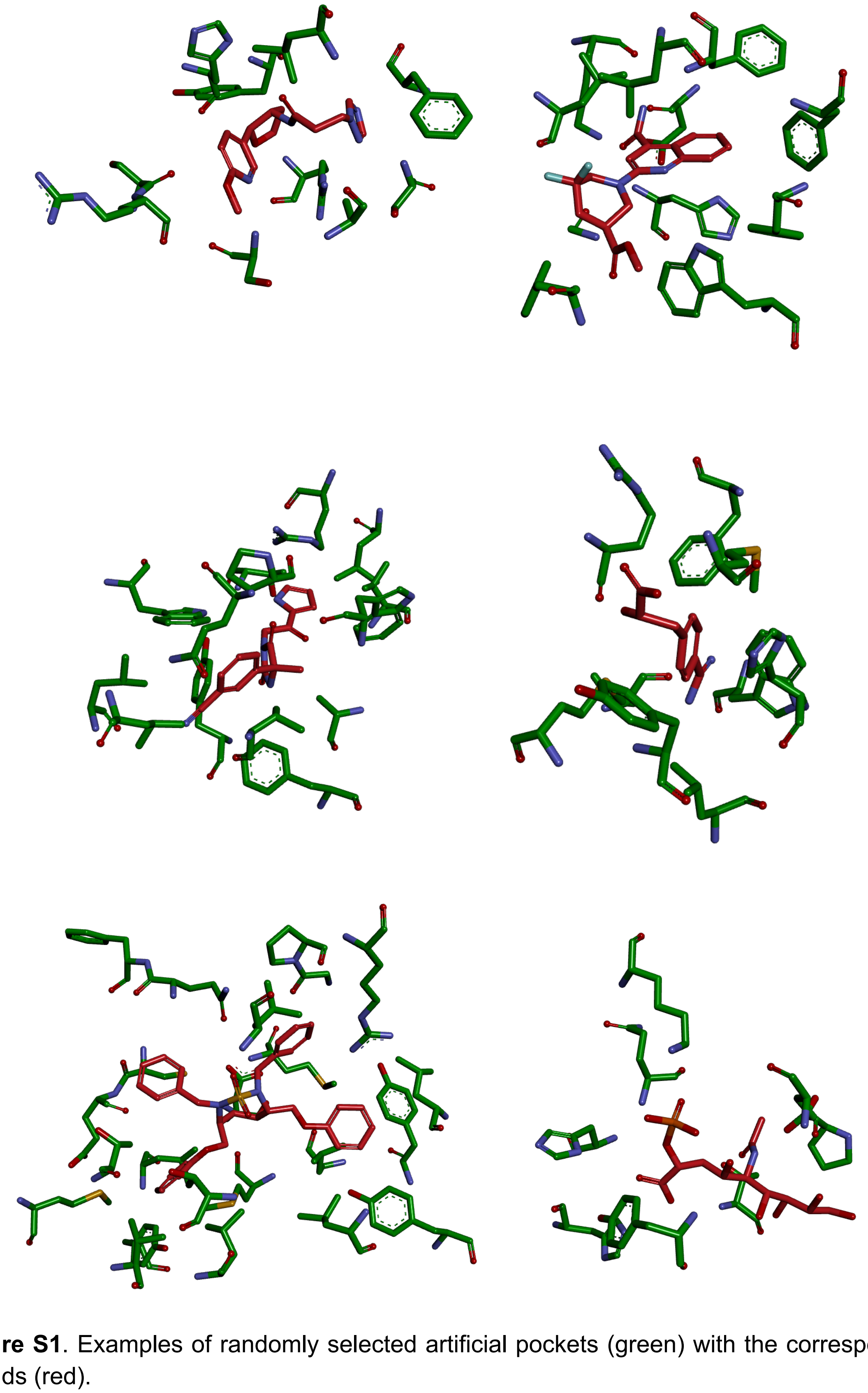

**Figure S1**. Examples of randomly selected artificial pockets (green) with the corresponding ligands (red).

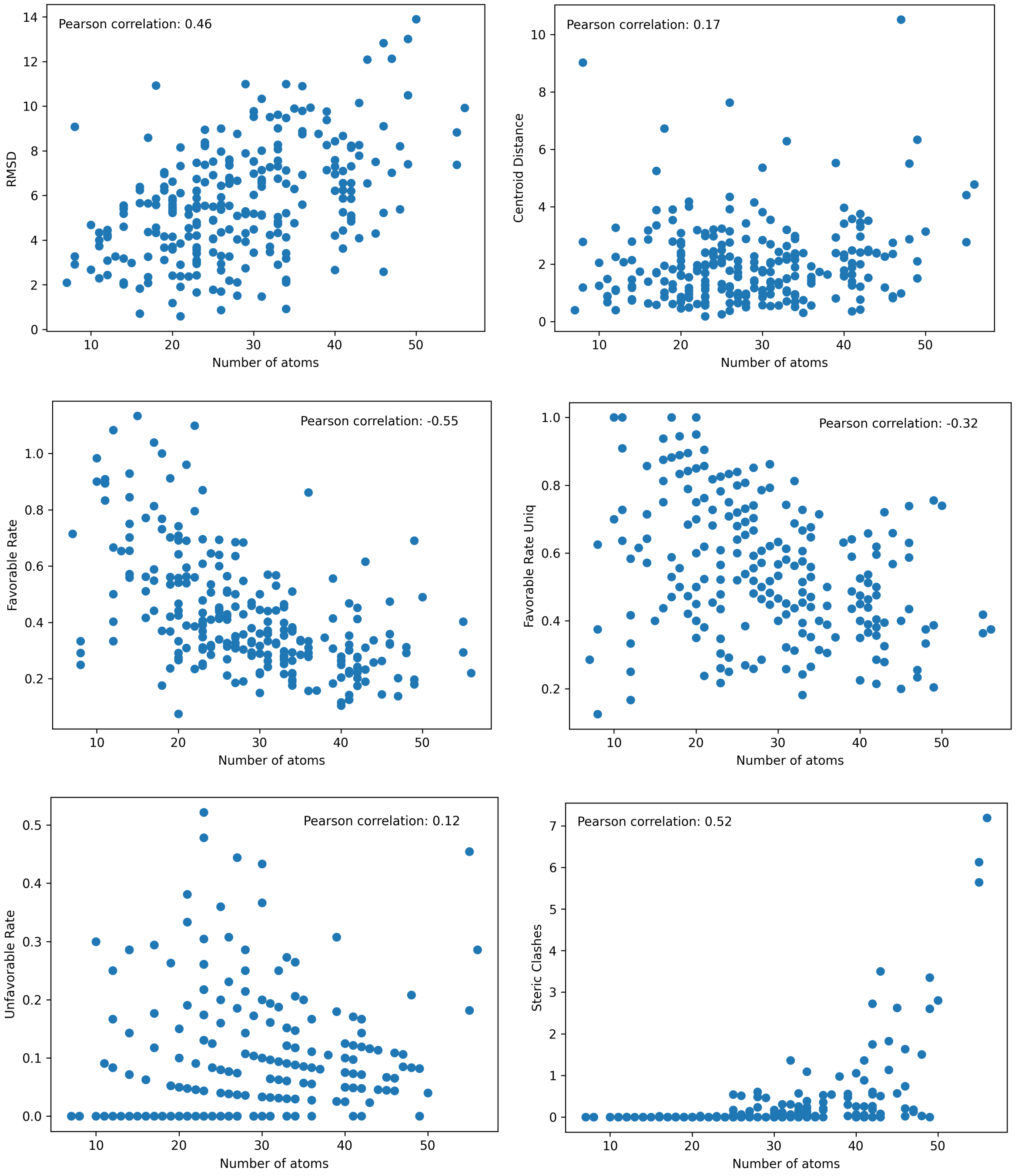


**Figure S2**. Correlation between the ligand heavy atom count and different evaluation metrics. The results are reported for the top-1 sample out of 40 predicted in the PDBbind test data.

**Table S1.** Characterisation of atoms in PDB residues.

| Residue Name | Atom Name | Hybridization | N Heavy Neighbors | Is Backbone | Is Hydrophobe | Is H Donor | Is Weak H Donor | Is H Acceptor | Is Positive | Is Negative | Is Aromatic |
|---|---|---|---|---|---|---|---|---|---|---|---|
| ALA | C | SP2 | 3 | 1 | 0 | 0 | 0 | 0 | 0 | 0 | 0 |
| ALA | CA | SP3 | 3 | 1 | 0 | 0 | 1 | 0 | 0 | 0 | 0 |
| ALA | CB | SP3 | 1 | 0 | 1 | 0 | 1 | 0 | 0 | 0 | 0 |
| ALA | N | SP2 | 2 | 1 | 0 | 1 | 0 | 0 | 0 | 0 | 0 |
| ALA | O | SP2 | 1 | 1 | 0 | 0 | 0 | 1 | 0 | 0 | 0 |
| ALA | OXT | SP2 | 1 | 1 | 0 | 0 | 0 | 1 | 0 | 1 | 0 |
| ARG | C | SP2 | 3 | 1 | 0 | 0 | 0 | 0 | 0 | 0 | 0 |
| ARG | CA | SP3 | 3 | 1 | 0 | 0 | 1 | 0 | 0 | 0 | 0 |
| ARG | CB | SP3 | 2 | 0 | 1 | 0 | 1 | 0 | 0 | 0 | 0 |
| ARG | CD | SP3 | 2 | 0 | 0 | 0 | 1 | 0 | 0 | 0 | 0 |
| ARG | CG | SP3 | 2 | 0 | 1 | 0 | 1 | 0 | 0 | 0 | 0 |
| ARG | CZ | SP2 | 3 | 0 | 0 | 0 | 0 | 0 | 0 | 0 | 0 |
| ARG | N | SP2 | 2 | 1 | 0 | 1 | 0 | 0 | 0 | 0 | 0 |
| ARG | NE | SP2 | 2 | 0 | 0 | 1 | 0 | 0 | 1 | 0 | 0 |
| ARG | NH1 | SP2 | 1 | 0 | 0 | 1 | 0 | 0 | 1 | 0 | 0 |
| ARG | NH2 | SP2 | 1 | 0 | 0 | 1 | 0 | 0 | 1 | 0 | 0 |
| ARG | O | SP2 | 1 | 1 | 0 | 0 | 0 | 1 | 0 | 0 | 0 |
| ARG | OXT | SP2 | 1 | 1 | 0 | 0 | 0 | 1 | 0 | 1 | 0 |
| ASN | C | SP2 | 3 | 1 | 0 | 0 | 0 | 0 | 0 | 0 | 0 |
| ASN | CA | SP3 | 3 | 1 | 0 | 0 | 1 | 0 | 0 | 0 | 0 |
| ASN | CB | SP3 | 2 | 0 | 1 | 0 | 1 | 0 | 0 | 0 | 0 |
| ASN | CG | SP2 | 3 | 0 | 0 | 0 | 0 | 0 | 0 | 0 | 0 |
| ASN | N | SP2 | 2 | 1 | 0 | 1 | 0 | 0 | 0 | 0 | 0 |
| ASN | ND2 | SP2 | 1 | 0 | 0 | 1 | 0 | 0 | 0 | 0 | 0 |
| ASN | O | SP2 | 1 | 1 | 0 | 0 | 0 | 1 | 0 | 0 | 0 |
| ASN | OD1 | SP2 | 1 | 0 | 0 | 0 | 0 | 1 | 0 | 0 | 0 |
| ASN | OXT | SP2 | 1 | 1 | 0 | 0 | 0 | 1 | 0 | 1 | 0 |
| ASP | C | SP2 | 3 | 1 | 0 | 0 | 0 | 0 | 0 | 0 | 0 |
| ASP | CA | SP3 | 3 | 1 | 0 | 0 | 1 | 0 | 0 | 0 | 0 |
| ASP | CB | SP3 | 2 | 0 | 1 | 0 | 1 | 0 | 0 | 0 | 0 |
| ASP | CG | SP2 | 3 | 0 | 0 | 0 | 0 | 0 | 0 | 0 | 0 |
| ASP | N | SP2 | 2 | 1 | 0 | 1 | 0 | 0 | 0 | 0 | 0 |
| ASP | O | SP2 | 1 | 1 | 0 | 0 | 0 | 1 | 0 | 0 | 0 |
| ASP | OD1 | SP2 | 1 | 0 | 0 | 0 | 0 | 1 | 0 | 1 | 0 |

| | | | | | | | | | | | |
|---|---|---|---|---|---|---|---|---|---|---|---|
| ASP | OD2 | SP2 | 1 | 0 | 0 | 0 | 0 | 1 | 0 | 1 | 0 |
| ASP | OXT | SP2 | 1 | 1 | 0 | 0 | 0 | 1 | 0 | 1 | 0 |
| CYS | C | SP2 | 3 | 1 | 0 | 0 | 0 | 0 | 0 | 0 | 0 |
| CYS | CA | SP3 | 3 | 1 | 0 | 0 | 1 | 0 | 0 | 0 | 0 |
| CYS | CB | SP3 | 2 | 0 | 1 | 0 | 1 | 0 | 0 | 0 | 0 |
| CYS | N | SP2 | 2 | 1 | 0 | 1 | 0 | 0 | 0 | 0 | 0 |
| CYS | O | SP2 | 1 | 1 | 0 | 0 | 0 | 1 | 0 | 0 | 0 |
| CYS | OXT | SP2 | 1 | 1 | 0 | 0 | 0 | 1 | 0 | 1 | 0 |
| CYS | SG | SP3 | 1 | 0 | 0 | 0 | 1 | 0 | 0 | 0 | 0 |
| GLN | C | SP2 | 3 | 1 | 0 | 0 | 0 | 0 | 0 | 0 | 0 |
| GLN | CA | SP3 | 3 | 1 | 0 | 0 | 1 | 0 | 0 | 0 | 0 |
| GLN | CB | SP3 | 2 | 0 | 1 | 0 | 1 | 0 | 0 | 0 | 0 |
| GLN | CD | SP2 | 3 | 0 | 0 | 0 | 0 | 0 | 0 | 0 | 0 |
| GLN | CG | SP3 | 2 | 0 | 1 | 0 | 1 | 0 | 0 | 0 | 0 |
| GLN | N | SP2 | 2 | 1 | 0 | 1 | 0 | 0 | 0 | 0 | 0 |
| GLN | NE2 | SP2 | 1 | 0 | 0 | 1 | 0 | 0 | 0 | 0 | 0 |
| GLN | O | SP2 | 1 | 1 | 0 | 0 | 0 | 1 | 0 | 0 | 0 |
| GLN | OE1 | SP2 | 1 | 0 | 0 | 0 | 0 | 1 | 0 | 0 | 0 |
| GLN | OXT | SP2 | 1 | 1 | 0 | 0 | 0 | 1 | 0 | 1 | 0 |
| GLU | C | SP2 | 3 | 1 | 0 | 0 | 0 | 0 | 0 | 0 | 0 |
| GLU | CA | SP3 | 3 | 1 | 0 | 0 | 1 | 0 | 0 | 0 | 0 |
| GLU | CB | SP3 | 2 | 0 | 1 | 0 | 1 | 0 | 0 | 0 | 0 |
| GLU | CD | SP2 | 3 | 0 | 0 | 0 | 0 | 0 | 0 | 0 | 0 |
| GLU | CG | SP3 | 2 | 0 | 1 | 0 | 1 | 0 | 0 | 0 | 0 |
| GLU | N | SP2 | 2 | 1 | 0 | 1 | 0 | 0 | 0 | 0 | 0 |
| GLU | O | SP2 | 1 | 1 | 0 | 0 | 0 | 1 | 0 | 0 | 0 |
| GLU | OE1 | SP2 | 1 | 0 | 0 | 0 | 0 | 1 | 0 | 1 | 0 |
| GLU | OE2 | SP2 | 1 | 0 | 0 | 0 | 0 | 1 | 0 | 1 | 0 |
| GLU | OXT | SP2 | 1 | 1 | 0 | 0 | 0 | 1 | 0 | 1 | 0 |
| GLY | C | SP2 | 3 | 1 | 0 | 0 | 0 | 0 | 0 | 0 | 0 |
| GLY | CA | SP3 | 2 | 1 | 0 | 0 | 1 | 0 | 0 | 0 | 0 |
| GLY | N | SP2 | 2 | 1 | 0 | 1 | 0 | 0 | 0 | 0 | 0 |
| GLY | O | SP2 | 1 | 1 | 0 | 0 | 0 | 1 | 0 | 0 | 0 |
| GLY | OXT | SP2 | 1 | 1 | 0 | 0 | 0 | 1 | 0 | 1 | 0 |
| HIS | C | SP2 | 3 | 1 | 0 | 0 | 0 | 0 | 0 | 0 | 0 |
| HIS | CA | SP3 | 3 | 1 | 0 | 0 | 1 | 0 | 0 | 0 | 0 |
| HIS | CB | SP3 | 2 | 0 | 1 | 0 | 1 | 0 | 0 | 0 | 0 |

| | | | | | | | | | | | |
|---|---|---|---|---|---|---|---|---|---|---|---|
| HIS | CD2 | SP2 | 2 | 0 | 0 | 0 | 0 | 0 | 0 | 0 | 1 |
| HIS | CE1 | SP2 | 2 | 0 | 0 | 0 | 0 | 0 | 0 | 0 | 1 |
| HIS | CG | SP2 | 3 | 0 | 0 | 0 | 0 | 0 | 0 | 0 | 1 |
| HIS | N | SP2 | 2 | 1 | 0 | 1 | 0 | 0 | 0 | 0 | 0 |
| HIS | ND1 | SP2 | 2 | 0 | 0 | 1 | 0 | 1 | 1 | 0 | 1 |
| HIS | NE2 | SP2 | 2 | 0 | 0 | 1 | 0 | 1 | 1 | 0 | 1 |
| HIS | O | SP2 | 1 | 1 | 0 | 0 | 0 | 1 | 0 | 0 | 0 |
| HIS | OXT | SP2 | 1 | 1 | 0 | 0 | 0 | 1 | 0 | 1 | 0 |
| ILE | C | SP2 | 3 | 1 | 0 | 0 | 0 | 0 | 0 | 0 | 0 |
| ILE | CA | SP3 | 3 | 1 | 0 | 0 | 1 | 0 | 0 | 0 | 0 |
| ILE | CB | SP3 | 3 | 0 | 1 | 0 | 1 | 0 | 0 | 0 | 0 |
| ILE | CD1 | SP3 | 1 | 0 | 1 | 0 | 1 | 0 | 0 | 0 | 0 |
| ILE | CG1 | SP3 | 2 | 0 | 1 | 0 | 1 | 0 | 0 | 0 | 0 |
| ILE | CG2 | SP3 | 1 | 0 | 1 | 0 | 1 | 0 | 0 | 0 | 0 |
| ILE | N | SP2 | 2 | 1 | 0 | 1 | 0 | 0 | 0 | 0 | 0 |
| ILE | O | SP2 | 1 | 1 | 0 | 0 | 0 | 1 | 0 | 0 | 0 |
| ILE | OXT | SP2 | 1 | 1 | 0 | 0 | 0 | 1 | 0 | 1 | 0 |
| LEU | C | SP2 | 3 | 1 | 0 | 0 | 0 | 0 | 0 | 0 | 0 |
| LEU | CA | SP3 | 3 | 1 | 0 | 0 | 1 | 0 | 0 | 0 | 0 |
| LEU | CB | SP3 | 2 | 0 | 1 | 0 | 1 | 0 | 0 | 0 | 0 |
| LEU | CD1 | SP3 | 1 | 0 | 1 | 0 | 1 | 0 | 0 | 0 | 0 |
| LEU | CD2 | SP3 | 1 | 0 | 1 | 0 | 1 | 0 | 0 | 0 | 0 |
| LEU | CG | SP3 | 3 | 0 | 1 | 0 | 1 | 0 | 0 | 0 | 0 |
| LEU | N | SP2 | 2 | 1 | 0 | 1 | 0 | 0 | 0 | 0 | 0 |
| LEU | O | SP2 | 1 | 1 | 0 | 0 | 0 | 1 | 0 | 0 | 0 |
| LEU | OXT | SP2 | 1 | 1 | 0 | 0 | 0 | 1 | 0 | 1 | 0 |
| LYS | C | SP2 | 3 | 1 | 0 | 0 | 0 | 0 | 0 | 0 | 0 |
| LYS | CA | SP3 | 3 | 1 | 0 | 0 | 1 | 0 | 0 | 0 | 0 |
| LYS | CB | SP3 | 2 | 0 | 1 | 0 | 1 | 0 | 0 | 0 | 0 |
| LYS | CD | SP3 | 2 | 0 | 1 | 0 | 1 | 0 | 0 | 0 | 0 |
| LYS | CE | SP3 | 2 | 0 | 0 | 0 | 1 | 0 | 0 | 0 | 0 |
| LYS | CG | SP3 | 2 | 0 | 1 | 0 | 1 | 0 | 0 | 0 | 0 |
| LYS | N | SP2 | 2 | 1 | 0 | 1 | 0 | 0 | 0 | 0 | 0 |
| LYS | NZ | SP3 | 1 | 0 | 0 | 1 | 0 | 0 | 1 | 0 | 0 |
| LYS | O | SP2 | 1 | 1 | 0 | 0 | 0 | 1 | 0 | 0 | 0 |
| LYS | OXT | SP2 | 1 | 1 | 0 | 0 | 0 | 1 | 0 | 1 | 0 |
| MET | C | SP2 | 3 | 1 | 0 | 0 | 0 | 0 | 0 | 0 | 0 |

| | | | | | | | | | | | |
|---|---|---|---|---|---|---|---|---|---|---|---|
| MET | CA | SP3 | 3 | 1 | 0 | 0 | 1 | 0 | 0 | 0 | 0 |
| MET | CB | SP3 | 2 | 0 | 1 | 0 | 1 | 0 | 0 | 0 | 0 |
| MET | CE | SP3 | 1 | 0 | 1 | 0 | 1 | 0 | 0 | 0 | 0 |
| MET | CG | SP3 | 2 | 0 | 1 | 0 | 1 | 0 | 0 | 0 | 0 |
| MET | N | SP2 | 2 | 1 | 0 | 1 | 0 | 0 | 0 | 0 | 0 |
| MET | O | SP2 | 1 | 1 | 0 | 0 | 0 | 1 | 0 | 0 | 0 |
| MET | OXT | SP2 | 1 | 1 | 0 | 0 | 0 | 1 | 0 | 1 | 0 |
| MET | SD | SP3 | 2 | 0 | 1 | 0 | 0 | 0 | 0 | 0 | 0 |
| PHE | C | SP2 | 3 | 1 | 0 | 0 | 0 | 0 | 0 | 0 | 0 |
| PHE | CA | SP3 | 3 | 1 | 0 | 0 | 1 | 0 | 0 | 0 | 0 |
| PHE | CB | SP3 | 2 | 0 | 1 | 0 | 1 | 0 | 0 | 0 | 0 |
| PHE | CD1 | SP2 | 2 | 0 | 1 | 0 | 1 | 0 | 0 | 0 | 1 |
| PHE | CD2 | SP2 | 2 | 0 | 1 | 0 | 1 | 0 | 0 | 0 | 1 |
| PHE | CE1 | SP2 | 2 | 0 | 1 | 0 | 1 | 0 | 0 | 0 | 1 |
| PHE | CE2 | SP2 | 2 | 0 | 1 | 0 | 1 | 0 | 0 | 0 | 1 |
| PHE | CG | SP2 | 3 | 0 | 1 | 0 | 0 | 0 | 0 | 0 | 1 |
| PHE | CZ | SP2 | 2 | 0 | 1 | 0 | 1 | 0 | 0 | 0 | 1 |
| PHE | N | SP2 | 2 | 1 | 0 | 1 | 0 | 0 | 0 | 0 | 0 |
| PHE | O | SP2 | 1 | 1 | 0 | 0 | 0 | 1 | 0 | 0 | 0 |
| PHE | OXT | SP2 | 1 | 1 | 0 | 0 | 0 | 1 | 0 | 1 | 0 |
| PRO | C | SP2 | 3 | 1 | 0 | 0 | 0 | 0 | 0 | 0 | 0 |
| PRO | CA | SP3 | 3 | 1 | 0 | 0 | 1 | 0 | 0 | 0 | 0 |
| PRO | CB | SP3 | 2 | 0 | 1 | 0 | 1 | 0 | 0 | 0 | 0 |
| PRO | CD | SP3 | 2 | 0 | 0 | 0 | 1 | 0 | 0 | 0 | 0 |
| PRO | CG | SP3 | 2 | 0 | 1 | 0 | 1 | 0 | 0 | 0 | 0 |
| PRO | N | SP2 | 3 | 1 | 0 | 0 | 0 | 0 | 0 | 0 | 0 |
| PRO | O | SP2 | 1 | 1 | 0 | 0 | 0 | 1 | 0 | 0 | 0 |
| PRO | OXT | SP2 | 1 | 1 | 0 | 0 | 0 | 1 | 0 | 1 | 0 |
| SER | C | SP2 | 3 | 1 | 0 | 0 | 0 | 0 | 0 | 0 | 0 |
| SER | CA | SP3 | 3 | 1 | 0 | 0 | 1 | 0 | 0 | 0 | 0 |
| SER | CB | SP3 | 2 | 0 | 0 | 0 | 1 | 0 | 0 | 0 | 0 |
| SER | N | SP2 | 2 | 1 | 0 | 1 | 0 | 0 | 0 | 0 | 0 |
| SER | O | SP2 | 1 | 1 | 0 | 0 | 0 | 1 | 0 | 0 | 0 |
| SER | OG | SP3 | 1 | 0 | 0 | 1 | 0 | 1 | 0 | 0 | 0 |
| SER | OXT | SP2 | 1 | 1 | 0 | 0 | 0 | 1 | 0 | 1 | 0 |
| THR | C | SP2 | 3 | 1 | 0 | 0 | 0 | 0 | 0 | 0 | 0 |
| THR | CA | SP3 | 3 | 1 | 0 | 0 | 1 | 0 | 0 | 0 | 0 |

| | | | | | | | | | | | |
|---|---|---|---|---|---|---|---|---|---|---|---|
| THR | CB | SP3 | 3 | 0 | 0 | 0 | 1 | 0 | 0 | 0 | 0 |
| THR | CG2 | SP3 | 1 | 0 | 1 | 0 | 1 | 0 | 0 | 0 | 0 |
| THR | N | SP2 | 2 | 1 | 0 | 1 | 0 | 0 | 0 | 0 | 0 |
| THR | O | SP2 | 1 | 1 | 0 | 0 | 0 | 1 | 0 | 0 | 0 |
| THR | OG1 | SP3 | 1 | 0 | 0 | 1 | 0 | 1 | 0 | 0 | 0 |
| THR | OXT | SP2 | 1 | 1 | 0 | 0 | 0 | 1 | 0 | 1 | 0 |
| TRP | C | SP2 | 3 | 1 | 0 | 0 | 0 | 0 | 0 | 0 | 0 |
| TRP | CA | SP3 | 3 | 1 | 0 | 0 | 1 | 0 | 0 | 0 | 0 |
| TRP | CB | SP3 | 2 | 0 | 1 | 0 | 1 | 0 | 0 | 0 | 0 |
| TRP | CD1 | SP2 | 2 | 0 | 0 | 0 | 1 | 0 | 0 | 0 | 1 |
| TRP | CD2 | SP2 | 3 | 0 | 1 | 0 | 0 | 0 | 0 | 0 | 1 |
| TRP | CE2 | SP2 | 3 | 0 | 0 | 0 | 0 | 0 | 0 | 0 | 1 |
| TRP | CE3 | SP2 | 2 | 0 | 1 | 0 | 1 | 0 | 0 | 0 | 1 |
| TRP | CG | SP2 | 3 | 0 | 1 | 0 | 0 | 0 | 0 | 0 | 1 |
| TRP | CH2 | SP2 | 2 | 0 | 1 | 0 | 1 | 0 | 0 | 0 | 1 |
| TRP | CZ2 | SP2 | 2 | 0 | 1 | 0 | 1 | 0 | 0 | 0 | 1 |
| TRP | CZ3 | SP2 | 2 | 0 | 1 | 0 | 1 | 0 | 0 | 0 | 1 |
| TRP | N | SP2 | 2 | 1 | 0 | 1 | 0 | 0 | 0 | 0 | 0 |
| TRP | NE1 | SP2 | 2 | 0 | 0 | 1 | 0 | 0 | 0 | 0 | 1 |
| TRP | O | SP2 | 1 | 1 | 0 | 0 | 0 | 1 | 0 | 0 | 0 |
| TRP | OXT | SP2 | 1 | 1 | 0 | 0 | 0 | 1 | 0 | 1 | 0 |
| TYR | C | SP2 | 3 | 1 | 0 | 0 | 0 | 0 | 0 | 0 | 0 |
| TYR | CA | SP3 | 3 | 1 | 0 | 0 | 1 | 0 | 0 | 0 | 0 |
| TYR | CB | SP3 | 2 | 0 | 1 | 0 | 1 | 0 | 0 | 0 | 0 |
| TYR | CD1 | SP2 | 2 | 0 | 1 | 0 | 1 | 0 | 0 | 0 | 1 |
| TYR | CD2 | SP2 | 2 | 0 | 1 | 0 | 1 | 0 | 0 | 0 | 1 |
| TYR | CE1 | SP2 | 2 | 0 | 1 | 0 | 1 | 0 | 0 | 0 | 1 |
| TYR | CE2 | SP2 | 2 | 0 | 1 | 0 | 1 | 0 | 0 | 0 | 1 |
| TYR | CG | SP2 | 3 | 0 | 1 | 0 | 0 | 0 | 0 | 0 | 1 |
| TYR | CZ | SP2 | 3 | 0 | 0 | 0 | 0 | 0 | 0 | 0 | 1 |
| TYR | N | SP2 | 2 | 1 | 0 | 1 | 0 | 0 | 0 | 0 | 0 |
| TYR | O | SP2 | 1 | 1 | 0 | 0 | 0 | 1 | 0 | 0 | 0 |
| TYR | OH | SP2 | 1 | 0 | 0 | 1 | 0 | 1 | 0 | 0 | 0 |
| TYR | OXT | SP2 | 1 | 1 | 0 | 0 | 0 | 1 | 0 | 1 | 0 |
| VAL | C | SP2 | 3 | 1 | 0 | 0 | 0 | 0 | 0 | 0 | 0 |
| VAL | CA | SP3 | 3 | 1 | 0 | 0 | 1 | 0 | 0 | 0 | 0 |
| VAL | CB | SP3 | 3 | 0 | 1 | 0 | 1 | 0 | 0 | 0 | 0 |

| | | | | | | | | | | |
|---|---|---|---|---|---|---|---|---|---|---|
| VAL | CG1 | SP3 | 1 | 0 | 1 | 0 | 1 | 0 | 0 | 0 | 0 |
| VAL | CG2 | SP3 | 1 | 0 | 1 | 0 | 1 | 0 | 0 | 0 | 0 |
| VAL | N | SP2 | 2 | 1 | 0 | 1 | 0 | 0 | 0 | 0 | 0 |
| VAL | O | SP2 | 1 | 1 | 0 | 0 | 0 | 1 | 0 | 0 | 0 |
| VAL | OXT | SP2 | 1 | 1 | 0 | 0 | 0 | 1 | 0 | 1 | 0 |

**Table S2.** Probabilities of the ligand atom, aromatic ring or amide group participation in the non-covalent interactions in the PDBbind complexes.

| **Protein** | **Ligand** | **Fraction** | **N interactions** | **N interacting uniq ligand elements** | **N compatible uniq ligand elements** | **Ligand element interaction probability** | **Avg n of ligand Interactions per element** |
|---|---|---|---|---|---|---|---|
| Hydrophobe (C, S) | Hydrophobe (C, S) | 74.70% | 255262 | 81217 | 98390 | 0.83 | 3.14 |
| Hydrophobe (C, S) | Hydrophobe (Cl, Br, I) | 2.76% | 9423 | 1932 | 2043 | 0.95 | 4.88 |
| Hydrophobe (C, S) | Hydrophobe (F) | 0.90% | 3089 | 1339 | 1737 | 0.77 | 2.31 |
| Aromatic Ring | Aromatic Ring | 1.80% | 6143 | 4999 | 18463 | 0.27 | 1.23 |
| Amide Group | Aromatic Ring | 1.40% | 4775 | 3847 | 18463 | 0.21 | 1.24 |
| Aromatic Ring | Amide Group | 0.32% | 1087 | 930 | 6982 | 0.13 | 1.17 |
| Acceptor | Donor | 6.69% | 22861 | 14692 | 23068 | 0.64 | 1.56 |
| Donor | Acceptor | 8.70% | 29742 | 20889 | 45843 | 0.46 | 1.42 |
| Acceptor | Halogen (Cl, Br, I) | 0.22% | 748 | 670 | 2043 | 0.33 | 1.12 |
| Negative | Positive | 0.67% | 2274 | 1802 | 4206 | 0.43 | 1.26 |
| Positive | Negative | 1.68% | 5757 | 4024 | 8427 | 0.48 | 1.43 |
| Aromatic Ring | Positive | 0.16% | 546 | 429 | 4206 | 0.10 | 1.27 |
| Positive | Aromatic Ring | 0.61% | 2077 | 1507 | 18463 | 0.08 | 1.38 |

**Table S3.** Residues frequencies in the PDBbind complexes.

| Residue Name | N |
|---|---|
| LEU | 15946 |
| GLY | 14757 |
| VAL | 12227 |
| ASP | 11081 |
| ILE | 10659 |
| TYR | 10523 |
| PHE | 10180 |
| ALA | 9418 |
| SER | 9182 |
| GLU | 8370 |
| HIS | 7897 |
| THR | 7864 |
| ARG | 6948 |
| ASN | 5997 |
| MET | 5770 |
| LYS | 5657 |
| TRP | 5203 |
| GLN | 4654 |
| PRO | 4539 |
| CYS | 3618 |

**Table S4.** Non-covalent protein-ligand interactions and their parameters. The distances for aromatic rings and amide groups are measured relative to their centroids. θ refers to an angle between the normal of the aromatic/amide plane and the vector between the aromatic ring centroid and the cation. For the hydrogen and halogen bonds, D is the donor atom covalently bound to the hydrogen/halogen; H/X is the hydrogen/halogen atom; A is an acceptor atom; Y is a heavy covalently bound neighbor of the acceptor atom. Multiple H-A-Y and X-A-Y angles may exist when there are more than one Y atoms in the structure. In such cases, the smallest angle is chosen. The minimal distances of favorable interactions is limited by the steric overlap threshold. The exceptions are the hydrogen bonds that might be nearly as short as the covalent bonds.

| Type | Interaction | Orientation | Distance range, Å | Angles range (degrees) |
|---|---|---|---|---|
| **Favorable interactions** | | | | |
| Hydrophobic | (C, S) - (C, S, Cl, Br, I) | | < 4.5 | |
| Hydrophobic | (C, S) - (F) | | < 3.9 | |
| Hydrophobic | Aromatic - Aromatic | face-to-face (parallel) | < 4.5 | θ ∈ {-30,30} |
| Hydrophobic | Aromatic - Aromatic | edge-to-face (T-shaped) | < 5.5 | θ ∈ {60,120} |
| Hydrophobic | Aromatic - Amide | face-to-face (parallel) | < 4.5 | θ ∈ {-30,30} |
| Hydrophobic | Aromatic - Amide | edge-to-face (T-shaped) | < 5.0 | θ ∈ {60,120} |
| Hydrogen Bond | Donor - Acceptor | | {2.5,3.9} | ∠D-H-A ≥ 90 ∠H-A-Y ≥ 90 |
| Halogen Bond | Cl, Br, I - Acceptor | | < 4.0 | ∠D-X-A ≥ 120 ∠X-A-Y ≥ 75 |
| Electrostatic | Positive - Negative | | {3.4,4.5} | |
| Electrostatic | Positive - Aromatic | face-to-face (above plane) | {3.4,4.5} | θ ∈ {-40,40} |
| **Unfavorable interactions** | | | | |
| | Donor - Donor | | < 3.4 | |
| | Acceptor - Acceptor | | < 3.2 | |
| | Positive - Positive | | < 5.2 | |
| | Negative - Negative | | < 5.2 | |
